# Inferring Chromosome Radial Organization from Hi-C Data

**DOI:** 10.1101/863803

**Authors:** Priyojit Das, Tongye Shen, Rachel Patton McCord

## Abstract

**Background:** The nonrandom radial organization of eukaryotic chromosome territories (CTs) inside the nucleus plays an important role in nuclear functional compartmentalization. Increasingly, chromosome conformation capture (Hi-C) based approaches are being used to characterize the genome structure of many cell types and conditions. Computational methods to extract 3D arrangements of CTs from this type of pairwise contact data will thus increase our ability to analyze CT organization in a wider variety of biological situations.

**Results:** A number of full-scale polymer models have successfully reconstructed the 3D structure of chromosome territories from Hi-C. To supplement such methods, we explore alternative, direct, and less computationally intensive approaches to capture radial CT organization from Hi-C data. We show that we can infer relative chromosome ordering using PCA on a thresholded inter-chromosomal contact matrix. We simulate an ensemble of possible CT arrangements using a force-directed network layout algorithm and propose an approach to integrate additional chromosome properties into our predictions. Our CT radial organization predictions have a high correlation with microscopy imaging data for various cell nucleus geometries (lymphoblastoid, skin fibroblast, and breast epithelial cells), and we can capture previously documented changes in senescent and progeria cells.

**Conclusions:** Our analysis approaches provide rapid and modular approaches to screen for alterations in CT organization across widely available Hi-C data. We demonstrate which stages of the approach can extract meaningful information, and also describe limitations of pairwise contacts alone to predict absolute 3D positions.

## Background

The three dimensional (3D) structure of the human genome is composed of different structures at different length scales. At smaller length scales, nucleosome positions, loops, and topologically associating domains are the most salient features, followed by compartmentalization at a longer length scale [1]. At the largest scale of this genome organization, the 3D bodies of individual chromosomes arrange mostly as discrete entities, known as chromosome territories (CTs) [2]. The arrangement of all the CTs inside the nucleus with respect to nuclear center and periphery forms the higher-order genome architecture. This CT organization is nonrandom with respect to the nucleus periphery and can play important roles in different nuclear mechanisms ranging from DNA replication and gene expression to the processing of RNA [3]. Recently, it has been shown that chromosome territorial organization can also protect genome from deleterious rearrangements during DNA damage [4]. Alterations in CT organization can also be important during cell differentiation [5] and in different disease conditions. For example, in Hutchinson-Gilford progeria syndrome cells, chr18 shifts toward the nucleus interior as compared to its position in normal proliferating fibroblast cells [6], while certain gene-rich chromosomes localize near the periphery in blebs in progeria cells [7]. Alterations in CTs can influence the likelihood of chromosomal translocations. For example, during adipogenesis, chr12 and chr16 become spatially proximal, increasing the chance of translocation between those chromosomes, which is the driving event of liposarcoma tumorigenesis [5]. Recently, researchers have shown that in breast cancer, a gain in inter-chromosomal interactions for chrX correlates with its gene expression changes [8]. Approaches to characterize CT positions can thus further our understanding of the implications of CT arrangements in health and disease.

From careful microscopic measurements over the past several decades, largely using sequence specific probes in fluorescence in situ hybridization (FISH), principles of CT organization in certain cell types have been identified. It has been observed that CTs are often organized according to one of two different distributions: either a gene density or chromosome length based pattern [9, 10]. Opposing forces of gene activity and lamina associated domain (LAD) density on different chromosomes likely contribute to these different distributions [11, 12]. In general, LADs are likely to be repressive to gene activity and occur more frequently on gene poor chromosomes, which also tend to be longer than gene rich chromosomes [13]. The proliferation rate of cells and nuclear shape have also been implicated as factors influencing CT organization. For example, proliferating cells tend to follow a gene density-based organization compared to the length-based distribution in quiescent or senescent cells [14]. Further, the spherical human lymphocyte nucleus follows a gene density based organization, which is conserved across several related species [15, 16], while the chromosome size based distribution is more prevalent in ellipsoidal fibroblast nuclei [17]. Our understanding of the relationships between factors influencing CT distribution are limited, however, by the fact that relatively few different cell types have been characterized in depth by this type of microscopic analysis.

Thorough analysis of CT positions requires not only observing their average, but also the distribution of possible positions across the cell population. The position of each chromosome varies between cells in the population, even while following certain tendencies [18]. Throughout interphase the CT positions remain stable, but change from one generation to another during mitosis as the nuclear envelope is broken down and re-established [19, 20, 21, 22]. Improvements to microscopy experiments [23, 24] and image analysis methods make sampling this variation in CT position across the population increasingly feasible [25, 26, 27], yet such analyses of all CT positions in a large number of cells exist for only few cell types [28].

In contrast, genome wide chromosome conformation capture (Hi-C) approaches are being applied to characterize the 3D genome structure of rapidly increasing numbers of cell types and conditions for the past decade [1, 29, 30, 31]. This Hi-C technique and its variants capture a snapshot of pairwise chromosomal interactions ranging from specific enhancer-promoter interactions [32] to large scale inter-chromosomal interactions. Due to the effect of noise and high variability, the inter-chromosomal interactions have received less attention compared to their intra-chromosomal counterparts. In a past few years, with the improvement of experimental protocols, the effect of noise on inter-chromosomal contacts has been reduced and studies based on relevant inter-chromosomal interactions have been started to emerge [33, 34, 35, 36]. For example, genes corresponding to olfactory receptors from different chromosomes form specific inter-chromosomal contacts in mouse olfactory sensory neurons which strengthen upon differentiation from a progenitor cell [37]. In an another set of studies, by analyzing Hi-C inter-chromosomal contacts obtained from different malignant diseases, researchers have identified several novel chromosomal rearrangements [38, 39]. Though Hi-C does not directly capture the radial position of the CTs, approaches that infer 3D chromosome positioning information from Hi-C contact data provide a valuable supplement to microscopic data, greatly increasing the number of cell types for which CT positions can be analyzed. In this study, we explore a set of Hi-C analysis approaches focused on rapid and efficient prediction of CT radial organization, ranging from very simple direct calculations on the Hi-C contact matrix to a network model tuned by additional chromosome property information.

Many approaches have been developed to reconstruct the 3D folding of individual chromosomes and the genome from Hi-C data at different resolutions using restraint and polymer physics based approaches [40, 41]. Some models focus on detailed structures of local regions of chromatin rather than the whole genome, and for others, the primary focus is often to understand the mechanistic principles underlying the organization of interphase and metaphase genome rather than a prediction of CT arrangement [42, 43, 44, 45, 46, 47, 48]. But, some of the approaches have also explored the radial arrangement of CTs in their 3D models. An early restraint-based model of the 3D genome based on tethered chromosome conformation capture (TCC) data predicted CT positions that agreed with the major principles of lymphoblast genome organization characterized by microscopy [42]. In an another work, Hi-C maps were probabilistically deconvolutated into a population of single cell structures and the averaged radial position of the CTs predicted from those structures matched fairly well with microscopic measurements from a single cell type [49]. A series of coarse-grained polymer simulation based studies have been performed to characterize non-random organization of the chromosomes using gene activity and random and biological looping constraints [50, 51, 52]. In a recent polymer modeling based study, the researchers used Hi-C derived properties and a chromatin state based energy function to study the principles of the spatial as well as radial genome organization [53].

Because numerous arrangements can potentially be consistent with a set of Hi-C contacts, in many cases, whole genome 3D models generated from Hi-C also have to take into account external information in order to increase the accuracy of CT positioning predictions. For example, Stevens et al. combine imaging and Hi-C contacts on single cells to orient their 3D chromosome models [54]. Similarly, the Chrom3D algorithm combines lamina associated domain (LAD) data along with Hi-C data to capture spatial and radial organization of the chromosomes [55]. Di Stefano et al. used a steered molecular dynamics based approach to reconstruct diploid genome organization from Hi-C data [56]. Using significant Hi-C interactions as the constraints, the modeling technique was able to capture the preferential nuclear position of different genomic regions based on their gene density, lamina association and epigenetic marks.

To supplement this landscape of approaches to predict 3D genome conformations from Hi-C data, we have several specific goals in this study. We describe and test direct, computationally non-intensive analysis approaches that have the focused aim of inferring CT radial positions from Hi-C data rather than a full model of 3D chromosome folding. Specifically, we demonstrate that radial organization patterns can be inferred from PCA analysis of thresholded inter-chromosomal contact matrices and show the utility of a force-directed graph layout algorithm to infer the average and variation around the average CT positions. These approaches thus do not require the computational resources necessary to calculate high resolution polymer models, and could be used to screen for potentially important differences in CT organization across a wide range of cell types and conditions without creating detailed 3D models in each case. We further evaluate the strengths and limitations of Hi-C contact data when it is used toward the goal of inferring large scale 3D positions of chromosomes, and where additional reference information needs to be added to the contacts to generate reliable radial positioning information. We finally evaluate the different stages of our approach on a variety of cell types and conditions, comparing to a variety of published microscopic data and predicting additional details of CT organization where limited microscopy data exists.

## Results

### Thresholding contacts extracts meaningful chromosomal interaction patterns from Hi-C data

Hi-C experiments capture spatial genome organization by measuring the interaction frequency between different genomic fragments, yielding information about both intra-chromosomal and inter-chromosomal interactions. Inter-chromosomal interactions generally occur much less frequently than intra-chromosomal interactions, and true interactions are mixed in with noise that arises from random background ligation [57, 30]. Despite this inherent noise and sometimes low signal, the interactions between chromosomes also contain information that reflects the radial organization of the chromosome territories inside the nucleus. But, excluding contacts that may primarily reflect the background is important to prevent those contacts from masking the true signal. In order to extract strong interactions that are more likely to distinguish radial chromosome positions between cell types, we applied a thresholding technique to the genome-wide contact matrix (see **Methods** section). The thresholding cutoff *h*_*cut*_value was calculated by taking a certain percentile of all the genome-wide Hi-C interactions and then the interactions greater than this cutoff limit are considered as strong interactions. To determine the desired cutoff value, we compared a Hi-C matrix from a specific cell type with a corresponding simulated random ligation matrix (see **Methods**) for different values of *h*_*cut*_ ranging from the 5^*th*^ to 95^*th*^ percentile. We examined two different cell types having different nuclear shapes (Supplementary Table 1), since the nuclear shape has been observed to correlate to some extent with the non-random radial organization of the CTs. The human blood lymphoblastoid cell, GM12878, has a spherical nucleus and follows a gene density based radial CT organization [9]. On the other hand, BJ1-hTERT human skin fibroblast cell has an ellipsoidal nucleus, which shows a chromosome length based CT organization [17]. The whole genome Hi-C contact matrices were obtained from Sanders et al. [58].

For each *h*_*cut*_ value, the number of chromosomal contact bin pairs passing the threshold were summed between each pair of chromosomes to a single bin (Eqn. 4).

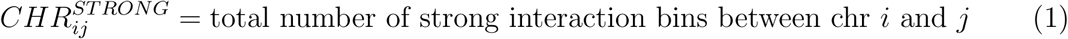

Then, Pearson’s correlation was applied to compare the whole chromosome pair interaction sums obtained from the original and corresponding random ligation Hi-C data. From Fig. 1a, it can be seen that with the increase in *h*_*cut*_ values from the 45^*th*^ to the 85^*th*^ percentile, the pairwise strong chromosomal interaction similarity between random and real data decreased rapidly and reached a stable value around the 90^*th*^ percentile. Similarly, we compared the strong chromosomal interaction sums with the pairwise product of the chromosome lengths to measure how much the number of interactions was primarily driven by chromosome length (i.e. two large chromosomes will have more interactions at random overall than two small chromosomes). This analysis produced a similar correlation trend as the random ligation effect comparison - strong interaction sums are no longer primarily explained by chromosome length at 90^*th*^ – 95^*th*^ percentile *h*_*cut*_ (Fig. 1a). Based on these two comparison results for both GM12878 and BJ1-hTERT, we chose the 95^*th*^ percentile as the final value of *h*_*cut*_ that leads to a minimized effect of chromosome length and random ligation for both cell types. While this optimal value was similar for two different datasets we considered, we note that Hi-C library complexity, read depth, and cis/trans ratio could affect the most appropriate *h*_*cut*_ value. As an alternative to this thresholding approach, we also explored the FitHiC [59] algorithm to extract significant chromosomal interactions from the Hi-C data. The analysis results obtained using those interactions are discussed in the following subsection.

**Figure 1.**
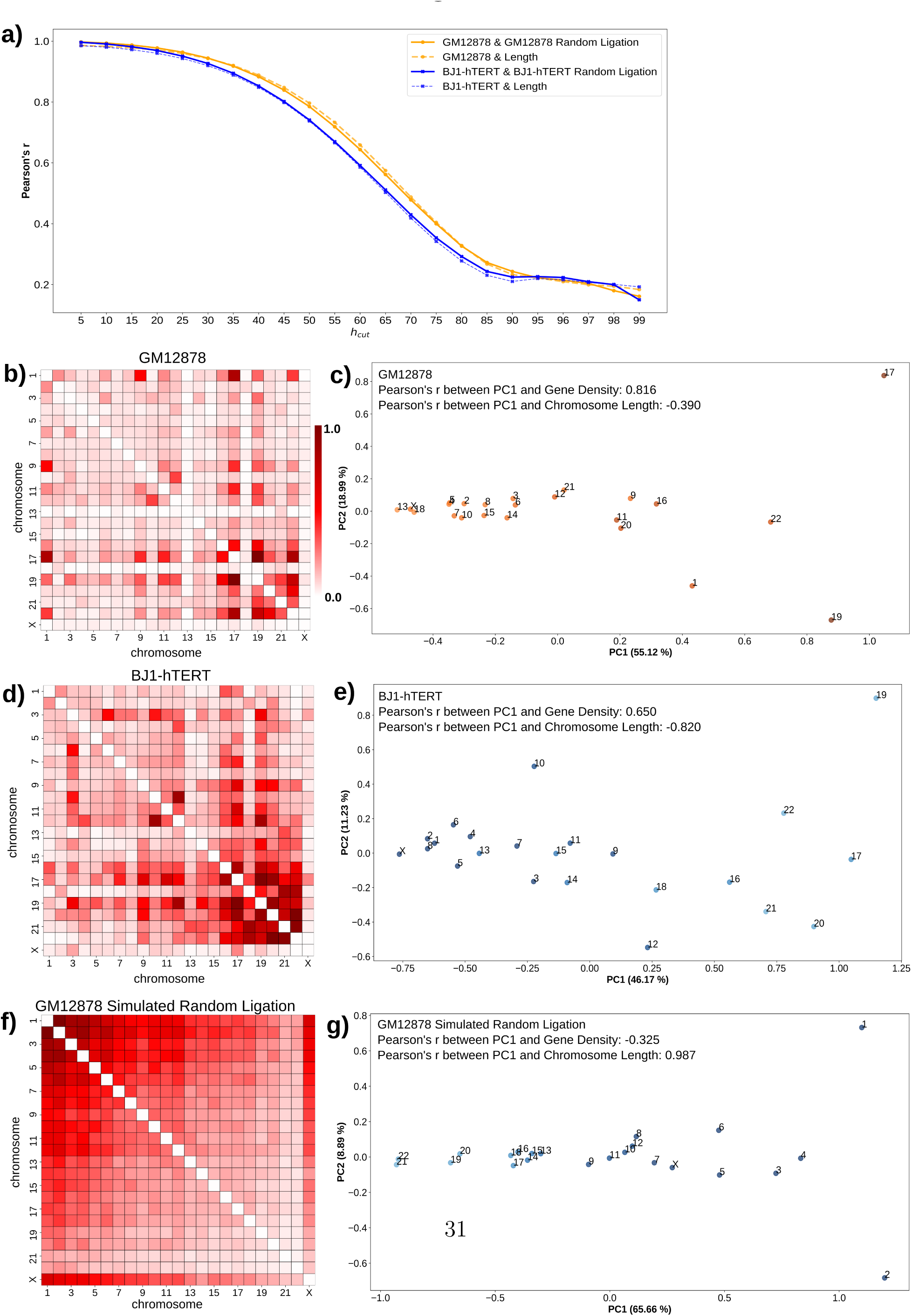
Determination of strong contact threshold and inference of radial CT distribution type. **a)** Correlation of pairwise chromosomal strong interaction pattern matrix obtained from original Hi-C with the pattern matrix from corresponding randomly ligated Hi-C (solid lines) and chromosome length (dotted lines) at different values of *h*_*cut*_. **b, d, f)** Pairwise inter-chromosomal strong interaction pattern matrix for GM12878 (**b**), BJ1-hTERT (**d**) and GM12878 simulated random ligation (**f**) Hi-C data respectively. Each entry of the matrix represents the total number of strong interacting bins between a pair of chromosomes. **c, e, g)** 2D PCA projection of the pairwise inter-chromosomal strong interaction pattern matrices obtained from GM12878 (**c**), BJ1-hTERT (**e**) and GM12878 simulated random ligation (**g**) Hi-C data respectively. Correlations of the PC1 projection with gene density and chromosome length (in bp) are shown. Chromosomes are color coded based on their gene density (orange) or length (blue). Dark color = high gene density/large chromosome size and light color = low gene density/short chromosome length.

### Radial chromosome ordering can be inferred from PCA on inter-chromosomal strong interaction pattern matrix

The pairwise chromosomal strong interaction pattern is not only able to distinguish the true biological interaction pattern from the random ligation, but also reveals distinct patterns specific to each cell type. By looking at the inter-chromosomal component of the pairwise strong interaction patterns (Eqn. 5) for GM12878 and

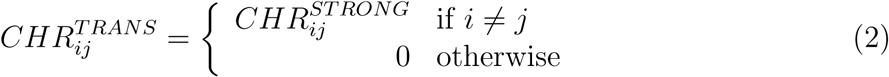

BJ1-hTERT represented in Fig. 1b and Fig. 1d respectively, we can clearly see that these two different cell types have distinct patterns - in BJ1-hTERT, the smaller chromosomes have higher strong inter-chromosomal interactions among themselves, whereas in GM12878 that pattern is dispersed. In order to capture the major interaction trends from those distinct patterns, we applied principal component analysis (PCA) to these matrices. This is similar to the application of PCA to contacts within chromosomes to detect A/B compartments or within proteins to detect structural domains [29, 60, 61]. Interestingly, we find that just as PCA can reveal the spatial segregation of domains within chromosomes, PCA on the pattern of strong contacts between all pairs of chromosomes can detect spatial segregation and relative ordering of chromosome territories. (Here we note that since the Hi-C matrix represents the average of the two copies of each chromosome, we use chromosome (chr) and chromosome territory (CT) interchangeably, noting that the CT positions will represent the average of the chromosome locations of each homolog across the cell population). Fig. 1c shows the 2D PCA projection of the pairwise inter-chromosomal strong interaction pattern matrix for the GM12878 cell. From this figure, it can be seen that chromosome 17, 19 and 22 and chromosome 13, 18 and X are on the opposite ends along the PC1 axis due to the high dissimilarity in their inter-chromosomal interaction patterns. In addition to that, this separation also correlates highly with the gene density based distribution of the chromosomes, as gene-rich chromosomes (e.g., chr19, chr17) are on the right extreme and gene-poor chromosomes (e.g., chr18, chr13) on the other extreme. This ordering thus corresponds to previous reports showing that lymphoblast cells with spherical nuclei tend to position their chromosomes in the nucleus in a gene density associated pattern [9]. Indeed, the PC1 values of the chromosomes show a higher absolute value correlation with their gene density than with their chromosome length. Next, we analyzed the pairwise inter-chromosomal significant interaction pattern matrix obtained using FitHiC for the GM12878 Hi-C data. We again see a higher PC1 correlation with gene density than chromosome length, but the correlation is much weaker (Supplementary Fig. 1b). This is not surprising given that the interaction pattern in the FitHiC chromosome pair interaction map is less distinct and more uniform across chromosomes (Supplementary Fig. 1a). Therefore, we choose to proceed with the strong interaction thresholding approach for our remaining analyses to detect the inherent radial arrangement of the CTs.

When we applied PCA to the BJ1-hTERT pairwise inter-chromosomal strong interaction pattern matrix, PC1 showed a different separation where chromosomes are ordered from left to right roughly according to decreasing length (Fig. 1e). The strong correlation between this ordering and chromosome length matches previous observations from fibroblast nuclei [17]. There is still some gene density correlation with this pattern, likely reflecting the complex picture of the radial CT organization in fibroblast nuclei. It has indeed been reported that while chromosomes are generally radially positioned by length in fibroblast nuclei, CT18 (gene poor and short) is still nearer to the nuclear envelope than CT19 (gene rich and short). When we applied PCA to the simulated random-ligation matrices generated from the GM12878 and BJ1-hTERT, we find that chromosomes are ordered along PC1 in a strong length-based distribution (Fig. 1g and Supplementary Fig. 2d). This is expected, since by chance longer chromosomes will have more random strong interactions than shorter chromosomes, as is evident in Fig. 1f.

The above results suggest that a simple application of PCA to the pairwise inter-chromosomal strong pattern matrix can reveal chromosome spatial distribution type, while also showing the limitations of this direct use of Hi-C contacts alone. Though PC1 can infer the radial CT distribution type, the direction of the ordering (whether inwards to outwards or outwards to inwards) is arbitrary and cannot be inferred based only this PCA result. Further, the random ligation result provides a caution that a length-based radial distribution would be detected even when no distinct interaction patterns are present.

To further explore the utility of such PCA ordering of CT positions, we next applied this analysis technique to Hi-C data from conditions which have been previously shown to exhibit large-scale rearrangement of chromosome territories. Specifically, in the premature aging disease Hutchinson-Gilford Progeria syndrome, it has been shown that chr18 moves to the nuclear interior compared to its peripheral location in normal proliferating fibroblasts, while chr10 shows the opposite trend [6, 62]. To check whether our analysis technique can also detect these CT reorganizations, we analyzed WI38-hTERT proliferating fibroblast Hi-C data [63] and Hi-C data from progeria patient fibroblasts at passage 19, when the cells are approaching senescence [64]. Fig. 2a and Fig. 2b show the 2D PCA projections of the pair-wise inter-chromosomal strong interaction pattern matrices obtained from the proliferating fibroblast and the progeria cells respectively. As mentioned above, while these projections reflect the underlying CT organization, inferring the directionality requires additional experimental information. There is evidence that chrX does not change its peripheral position in progeria cells compared to normal proliferating fibroblasts [6]. Indeed, in our analyses, chrX is positioned at a far extreme of PC1 in both progeria and proliferating fibroblasts, so we assigned the end of the PC1 axis near chrX as the periphery and the opposite end as the nucleus center, as seen in Fig. 2a and Fig. 2b. Now, if we examine the positions of chr18 in these projections, we can see that chr18 occupies a more internal position in the progeria cell (Fig. 2a and Fig. 2b). We can visualize the relative changes in chromosome positions by ordering the chromosomes based on the PC1 value in an increasing fashion from center to periphery in a radar plot (Fig. 2d). Here, we can see the internal shift of chr18 in progeria compared to the proliferating fibroblast. We did not find any significant change in the chr10 position in our analysis. However, we noticed that chr13 in progeria also moves to interior similar to chr18, which can be found in proliferating lamionpathy fibroblasts [62]. Progeria cells approaching senescence have a CT arrangement that to some extent matches other quiescent and senescent cells [6]. Therefore, next, we performed PCA ordering analysis on WI38-hTERT oncogene induced senescent cell Hi-C data [63]. When we compared the result with normal proliferating cells, we found chr10, chr13, chr18 and chrX radial rearrangements similar to the progeria cells (Fig. 2c and Fig. 2d bottom). Overall, this analysis shows that the strong inter-chromosomal interaction pattern from Hi-C has the potential to infer the underlying CT distribution within the nucleus.

**Figure 2.**
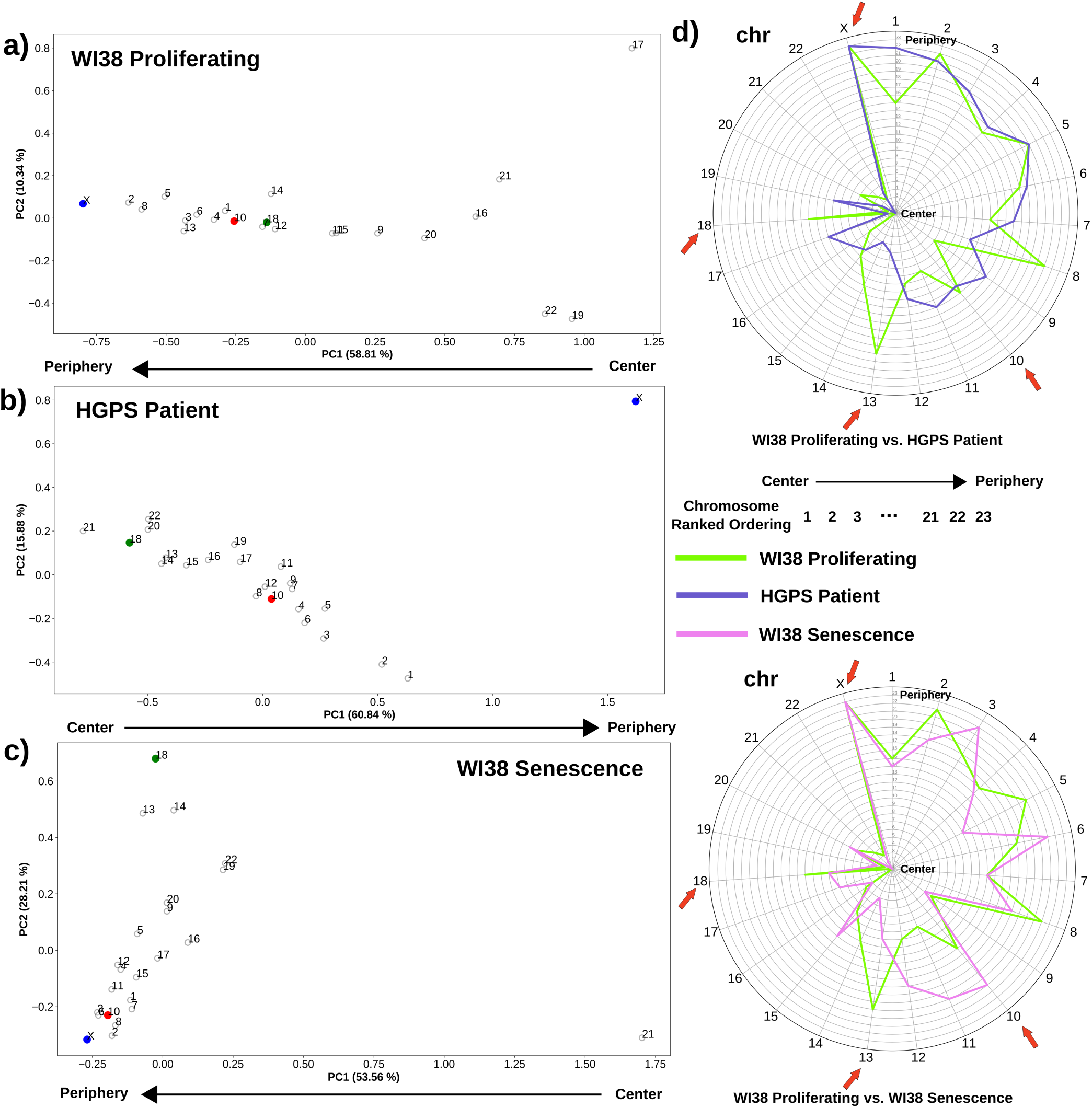
Rearrangement of radial CT position in progeria and senescent cells compared to proliferating fibroblasts. **a-c)** 2D PCA projection of the pairwise inter-chromosomal strong interaction pattern matrices obtained from WI38 proliferating (**a**), Hutchinson-Gilford progeria syndrome patient passage 19 (**b**) and WI38 senescent fibroblast (**c**) Hi-C data respectively. In progeria and senescent cells, chr10 (red color) and chr18 (green color) change their radial positions compared to the proliferating cells (chr10 change is not significant in progeria), while chrX maintains its peripheral position in all of the cases. **d)** Radar plot representing the change in the radial order of the CTs in progeria and senescence compared to the proliferating condition. Chromosome radial positions are shown as ordered (ranked) in an increasing fashion from the center to periphery. Salient chromosomes discussed in the text are highlighted with red arrows.

### Network modeling predicts probabilistic radial CT organization inside nucleus

Though applying a simple PCA approach to a pairwise inter-chromosomal strong interaction pattern matrix is surprisingly powerful at detecting relative chromosome ordering in different cell types, as shown above, it is limited to predicting a single position for each chromosome, and will predict ordering patterns even for data that is actually random noise. In reality, chromosome territory positions vary substantially within individual cells, even while following certain general trends. The positioning of CTs in individual cells changes from mother cell to daughter cell during mitosis [19, 20]. Thus, we sought an approach to model not only the average position, but the variability of chromosome positions in an ensemble of arrangements derived from Hi-C data. For this, we tested a network modeling approach. We used the strong interaction matrix obtained from the thresholding step to construct an undirected weighted graph in three-dimensional space using the 3D Fruchterman-Reingold (FR) force-based layout [65] algorithm. Each node of the graph represents a genomic region of fixed size and the weights are the contact frequencies. In such a graph layout, the nodes within each chromosome become clustered together due to high intra-chromosomal contacts while chromosomes that have higher inter-chromosomal contacts among themselves will end up close together in 3D space compared to the other chromosomes with less inter-chromosomal contacts. To model the variable CT arrangements possible within the average Hi-C data, 1000 independent runs of the FR algorithm were performed with different random initial configurations, which led to the formation of 1000 different 3D network graphs (see **Methods** section). Next, a minimum volume ellipsoid algorithm was used to fit a geometrical object around the models to serve as the nuclear periphery. From each of those models, we calculated the distance of the center of mass of each chromosome from the nucleus center (center of the geometrical fitted object).

When we examined the models, we found that they cluster into several different possible chromosome organization patterns (Supplementary Fig. 3). As we discovered with the PCA analysis, the Hi-C data alone cannot determine the absolute ordering from interior to periphery, so some clusters of models represent inverted patterns of organization. We also find some clusters of models that do not coincide with the length or gene density based distributions inferred from PCA CT ordering. These may represent local minima reached from certain initial conditions of the network model which do not reflect the true chromosome arrangements. Given these observations, we combine information from the previous PCA analysis with these network models to select the cluster of models for further analysis. After observing that the PCA analysis reports whether the cell type in general follows a length or gene density-based radial distribution, we use this information to select the cluster of models which has the highest absolute correlation between its mean CT distances and gene density or chromosome length (based on the inferred CT distribution type from the pairwise inter-chromosomal PCA transformation).

With the selected cluster of models (Fig. 3a and b), we next examined the predicted heterogeneity in chromosome territory positioning within each cell type (Fig. 3 and Supplementary Fig. 4). Fig. 3c shows the radial distance profiles of four example CTs - CT1, CT18, CT19 and CT20 obtained from GM1878 and BJ1-hTERT network model clusters. Among these CTs, CT18 (length 78.1 Mbp and gene density 3.4 genes / sequenced Mbp) and CT19 (length 59.1 Mbp and gene density 23.9 genes / sequenced Mbp) have comparable DNA content but have drastically different gene densities. The network modeling result shows a trend in which, in GM12878, CT18 has a peak near the periphery and CT19 has a more internal peak. On the other hand, in BJ1-hTERT, both CT18 and CT19 peaks are located toward the nuclear center with a similar distribution. The next contrasting pair is CT1 (length 249.3 Mb and gene density 8.70 genes / sequenced Mb) and CT20 (length 63.0 Mb and gene density 8.71 genes / sequenced Mb). They both have comparable gene densities but strikingly different lengths. Again based on modeling results, we can see that CT1 and CT20 have internal locations in GM12878, but in BJ1-hTERT CT1 occupies a much peripheral location. As expected given the overall gene density or length correlations of the model cluster, these mean positions are consistent with microscopy data regarding the different positioning of chr18, chr19, and chr1 in different cell types. Beyond mean position shifts, we also observe that the distributions of possible chromosome positions also match microscopic evidence for chromosome position variation in some respects. For example, in the models, CT4 in lymphoblasts is highly skewed toward peripheral positions and almost never observed near the center of the nucleus. This matches the distribution of positions of a gene located on chr4 measured by FISH [27] in which the gene was almost never observed in the interior 30 percent of the nuclear radius in any individual cell (Supplementary Fig. 5d). In contrast, the models predict that CT10 in fibroblasts can be found throughout the middle of the nucleus, but almost never at the extreme center or periphery (Supplementary Fig. 5c). This matches the measured chr10 distribution by FISH [66]. In some cases, however, the model-inferred CT distribution matches the mean position observed in microscopy, but not the distribution of positions. For example, the models locate chr19 near the nucleus center on average in both cell types we considered, as in microscopy data (Supplementary Fig. 5a and Supplementary Fig. 5b). However, in single cell images, chr19 is found over a broader range of positions [67], while our models predict a more tightly focused consistent positioning of chr19. This narrow predicted distribution likely stems from the highly specific interaction pattern for chr19, visible in the strong contact matrix and distinct PCA position of this chromosome in Fig. 1. Additionally, while chr13 has been observed by microscopy to be strongly skewed toward the nuclear periphery [66, 68], our model sometimes predicts a strongly peripheral distribution (GM12878) and sometimes a strongly internal distribution (BJ1-hTERT) (Supplementary Fig. 4). This occurs because there are relatively few strong inter-chromosomal interactions detected by chr13 in the Hi-C data, and the model cannot distinguish whether this means the chromosome is located on its own far to the center or far to the periphery. So, while our network model approach shows a strong ability to predict chromosome position variability for some chromosomes, in other cases, there are inherent limitations of what can be predicted from Hi-C contacts alone.

**Figure 3.**
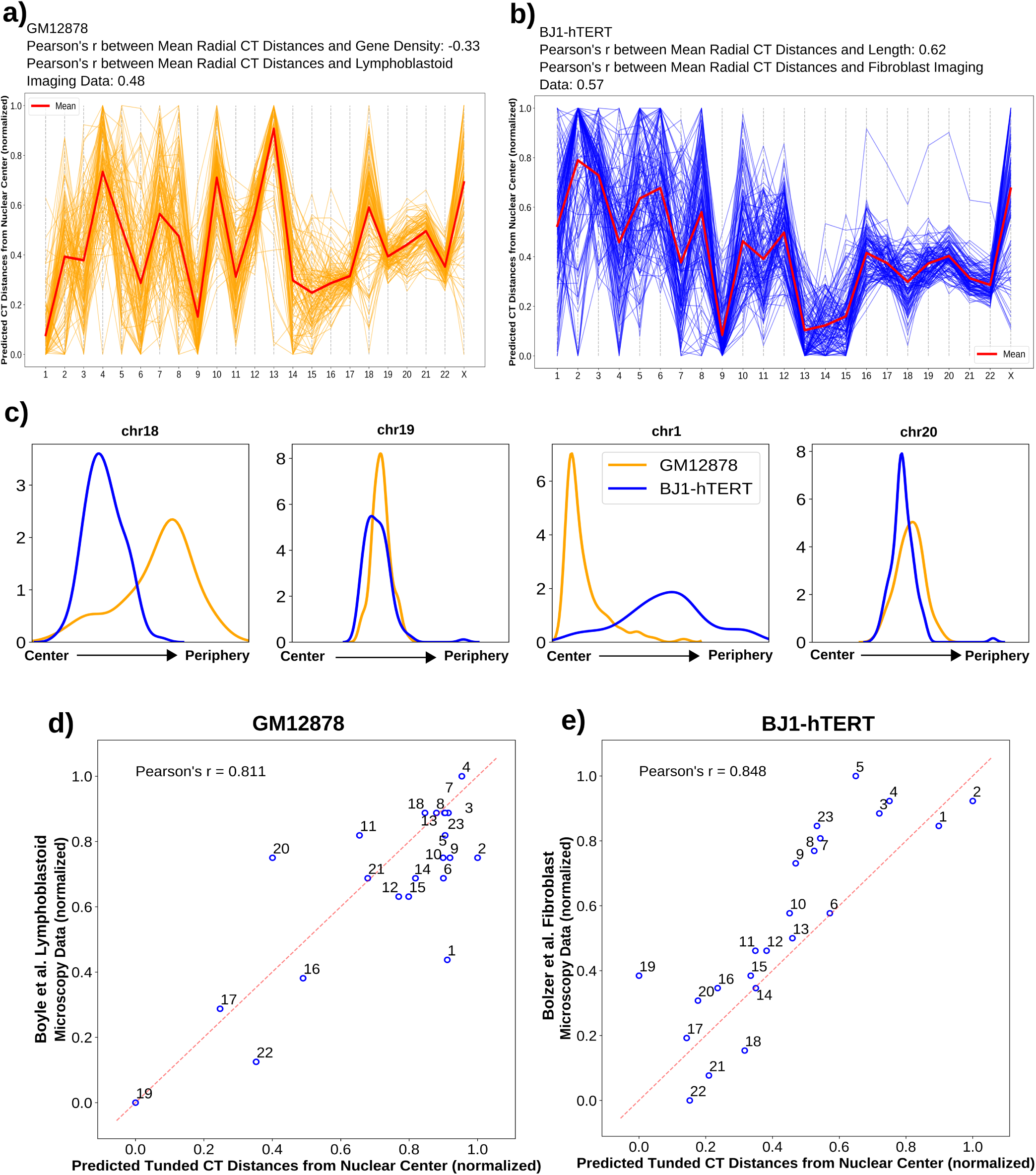
Network modeling generated distributions of CT radial positions and comparison of the predicted tuned result with microscopy data. **a-b)** Network modeling generated model clusters for both GM12878 (**a**) and BJ1-hTERT (**b**), selected based on respective inferred CT distribution types. **c)** The radial distance profiles of 4 CTs (chr18, chr19, chr1 and chr20) for both GM12878 and BJ1-hTERT obtained from the respective selected model clusters (# of models in the selected clusters: for GM12878 - 109, and BJ1-hTERT63122). **d)** Correlation between predicted tuned CT distance from GM12878 Hi-C data and lymphoblastoid microscopy imaging data. **e)** Correlation between predicted tuned CT distance from BJ1-hTERT Hi-C data and fibroblast microscopy imaging data.

In contrast to the original Hi-C contact maps, simulated random ligation maps produced network models that were much more variable and did not form tight clusters (Supplementary Fig. 2a and Supplementary Fig. 2e). This primarily happens due to non-specific pairwise inter-chromosomal interaction pattern driven by length which ultimately leads to the generation of wide variety of random network model configurations. Although, these simulated random ligation datasets show strong length based distribution of CTs from the PCA ordering, the pairwise inter-chromosomal strong interaction pattern is quite different from true length based distribution as shown by ellipsoidal cells. When we examine the distributions of chromosome positions predicted for these random models, most chromosomes showed broad indistinct distributions that did not vary based on the cell type the random matrix was derived from (Supplementary Fig. 6). This demonstrates that real Hi-C contacts contribute important information to our network modeling predicted radial distance distributions.

### Chromosome property-based tuning improves predicted consensus radial arrangement of CTs

Although the network modeling generated radial distribution of the CTs can reveal several interesting features of the CT organization that match with prior observations, the direct correlation with microscopy measured mean positions is only modest (Fig. 3a and Fig. 3b). These mean radial positions can further be fine-tuned by adding additional chromosomal properties explicitly. As it has already been shown that the radial organization of CTs depends on gene density and chromosome length to some extent, here, we added that effect to the averaged positions using weighted averaging and loess (locally estimated scatterplot smoothing) [69] techniques. This tuning procedure is described in detail in the **Methods** section, but the basic idea behind it involves several concepts. First, we incorporate information about the length and gene density of each chromosome explicitly. Second, going back to our earlier observation that the PC1 of the pairwise inter-chromosomal strong interaction pattern matrix can provide meaningful ordering information, we combine the network model output with PC1 ordering result obtained from selected model cluster through weighted averaging. Finally, in calculating weights for our averaging calculation, we take into account how much of the variance in the chromosome positioning pattern in the selected model cluster is captured by PC1 and PC2. We find this metric captures a major distinction between random and real Hi-C data. In the previous steps, random data will give some pattern, often highly chromosome length related, but we find that PC1 and PC2 explain a much lower percentage of the overall variance in the chromosome contact pattern for random data, while for real data these first two PCs capture most of the variance, indicating that the orderings along these PCs are highly meaningful. Thus, weighting by the variance explained by PC1 and PC2 will emphasize meaningful patterns over random patterns.

In order to evaluate the accuracy of the tuning procedure, we compared the tuned radial CT positions of GM12878 and BJ1-hTERT with corresponding microscopy imaging data, as shown in Fig. 3d and Fig. 3e. From Fig. 3d, it can be seen that, in GM12878, most of the CT positions correlate well with the experimentally obtained position with slight displacements, apart from the CT1 position, which moves outwards compared to the experimental data. Similarly, predicted CT distances for the BJ1-hTERT cell shows high similarity with the corresponding imaging data. For this particular cell, out of all the CTs, CT19 showed a higher amount of displacement towards the center in the predicted result. When the tuning technique was tested on simulated random ligation Hi-C data generated from GM12878 and BJ1-hTERT, it produced a far weaker correlation with the fibroblast imaging result in both cases (length based inferred CT distribution type) compared to the original Hi-C analysis results (Supplementary Fig. 2b and Supplementary Fig. 2f). Furthermore, we checked the consistency of the predicted tuned results between two Hi-C replicates of the BJ1-hTERT cell and found very little difference in chromosome positioning (Supplementary Fig. 7f). In case of GM12878, variation between replicates was higher, perhaps reflecting the larger variation in quality metrics between these two datasets (Supplementary Fig. 8f). However, a strength of employing the PCA ordering is revealed in comparing these replicates: the PCA ordering of the strong interaction matrix is robust to such variations in dataset quality (Supplementary Fig. 8d).

### Paternal and maternal homologs show mostly similar radial arrangement of CTs

Our modeling approach places only one copy of each chromosome in the radial organization network, and we assume that this represents the average of the two homolog positions. In any given cell, it is known from microscopy that the CT positions of homologs can be quite different [67], but we assume that both homologs would follow the same overall distribution of positions, and thus averaging their positions is reasonable. To test this assumption, we applied our approach to the deeply sequenced GM12878 Hi-C data from Rao et al. that can be mapped to maternal and paternal homologs based on allele-specific single nucleotide polymorphisms (SNPs) [70]. Once we had the genome wide Hi-C contact matrices for the paternal and maternal copies, thresholding was applied and pairwise inter-chromosomal strong interaction pattern matrix was analyzed using PCA transformation for each of them. Fig. 4c shows the 2D PCA projection of the pairwise inter-chromosomal strong interaction pattern matrix obtained from the paternal copy. By looking at this figure, we can clearly see that the paternal homologs are following a gene density driven distribution along the PC1 axis, which we also found for the the maternal homologs (Fig. 4d y-axis). In addition to that, the PC1 values from the paternal and maternal copies are highly correlated to each other and also with the PC1 values obtained from the previous GM12878 data (Fig. 4d and Fig. 4e). This suggests that the paternal and maternal homologs follow the similar radial distribution inside the nucleus, and that it is fair to infer a single average position for both homologs. We also applied the network modeling approach to the paternal and maternal copies individually and selected the model cluster for each of them whose mean CT distances has highest absolute correlation with the inferred CT distribution type, as above. By looking at the mean CT distances from the Fig. 4f and Fig. 4g, we can see that the mean radial CT distribution pattern between two copies are highly comparable with a single drastic exception in case of chr2, which shows an internal position in maternal copy and occupies a peripheral position in the paternal one. Again this can be explained by distinct patterns of inter-chromosomal strong interaction of chr2 paternal and maternal homologs. Finally, when we compared the radial distance profiles of the 4 CTs - CT1, CT18, CT19 and CT20 among the paternal, maternal and the standard GM12878, we found the density peaks in similar radial positions. However, in case of chr1, we found a strong narrow peak from the standard GM12878 data compared to the short broad peak obtained from both the paternal and maternal copies, which might arise due to the less specific inter-chromosomal interaction pattern of chr1 in both maternal and paternal copies compared to the interaction patterns of rest of the chromosomes in those copies. Overall, we observe that representing both homologs with an average predicted position is valid.

**Figure 4.**
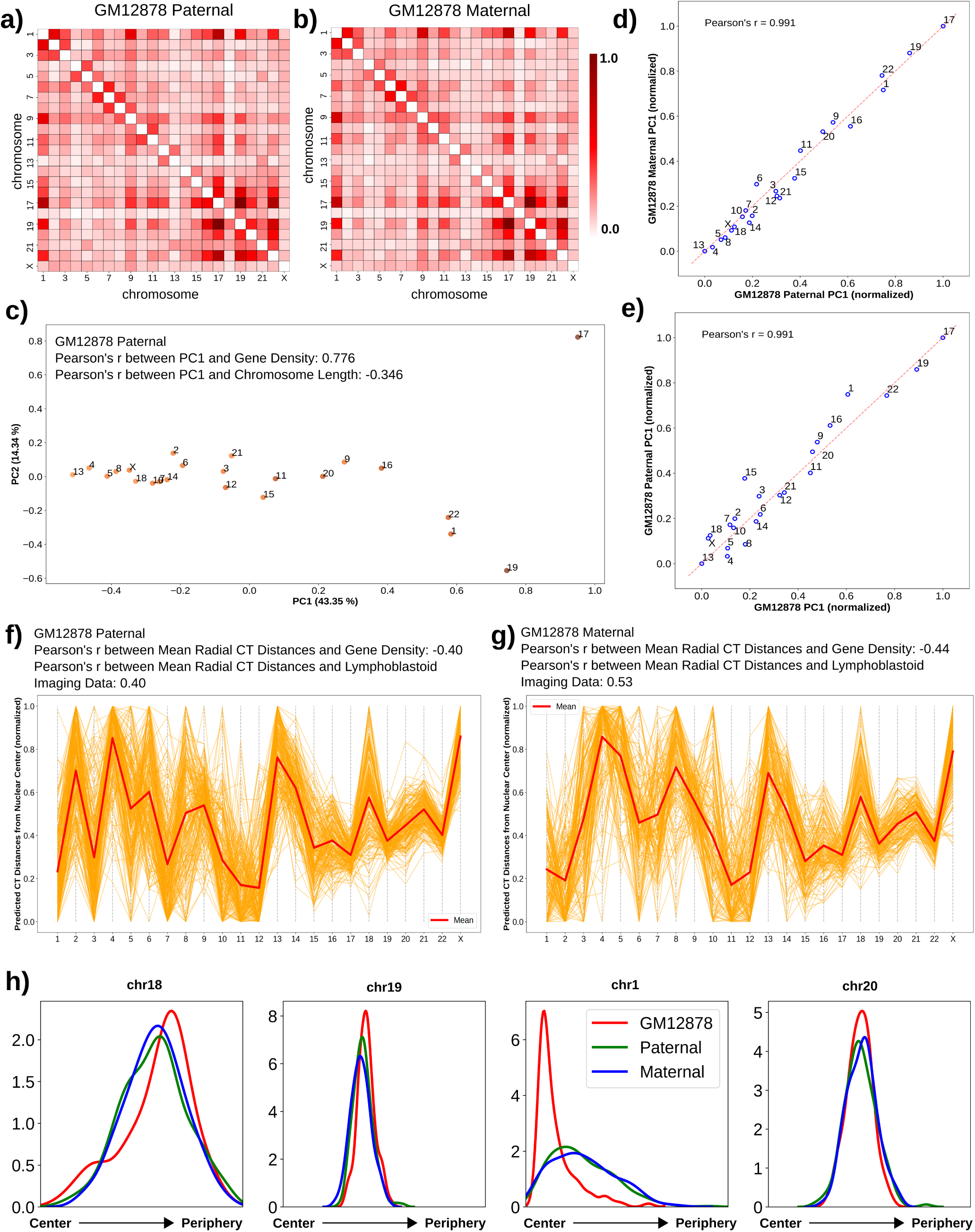
Paternal and maternal homologs follow a similar CT distribution pattern. **a-b)** Pairwise inter-chromosomal strong interaction pattern matrices for GM12878 paternal(**a**) and maternal (**b**) copies. **c)** 2D PCA projection of the pairwise inter-chromosomal strong interaction pattern matrix for GM12878 paternal copy. **d-e)** Correlation of GM12878 paternal copy PC1 values with maternal copy (**d**) and diploid averaged GM12878 (**e**) PC1 values obtained from the PCA transformation of the respective pairwise inter-chromosomal strong interaction pattern matrices. **f-g)** Network modeling generated model clusters for both GM12878 paternal (**f**) and maternal (**g**) copies, selected based on respective inferred CT distribution types. **h)** Predicted radial distance distributions of 4 CTs (chr18, chr19, chr1 and chr20) for GM12878 diploid averaged, paternal and maternal copies, obtained from the respective selected model clusters (# of models in the selected clusters: GM12878 diploid average - 109, paternal - 194, and maternal - 148).

### Radial CT organization in epithelial cells changes depending on their nuclear shape

After testing the performance of the analysis technique on the GM12878 and BJ1-hTERT cells, we applied our analysis to MCF10A non-tumorigenic breast epithelial cells. The Hi-C contact data of this cell was downloaded from Barutcu et al. [71] and binned at 2.5 Mb resolution. We obtained corresponding imaging data for 8 CTs (CT1, CT4, CT11, CT12, CT15, CT16, CT18 and CT21) of the MCF10A from Fritz et al. [8]. When we applied PCA to the pairwise inter-chromosomal strong interaction matrix, PC1 showed higher absolute correlation with chromosome length compared to gene density, as expected from experimental data (Fig. 5b), and corroborating the association of the length based distribution with ellipsoidal nucleus. Next, we used the network modeling approach to generate radial distance profiles of the 8 CTs and found the distributions match the ordering of CT positions reported in imaging data (Fig. 5c). We note that we predict that some chromosomes (chr4, chr15) have a much broader distribution than others (chr16, chr21), but this distribution is not reported for comparison in microscopy data. Finally, when the tuned CT positions were compared with the experimental data, a high correlation was obtained (Fig. 5d).

**Figure 5.**
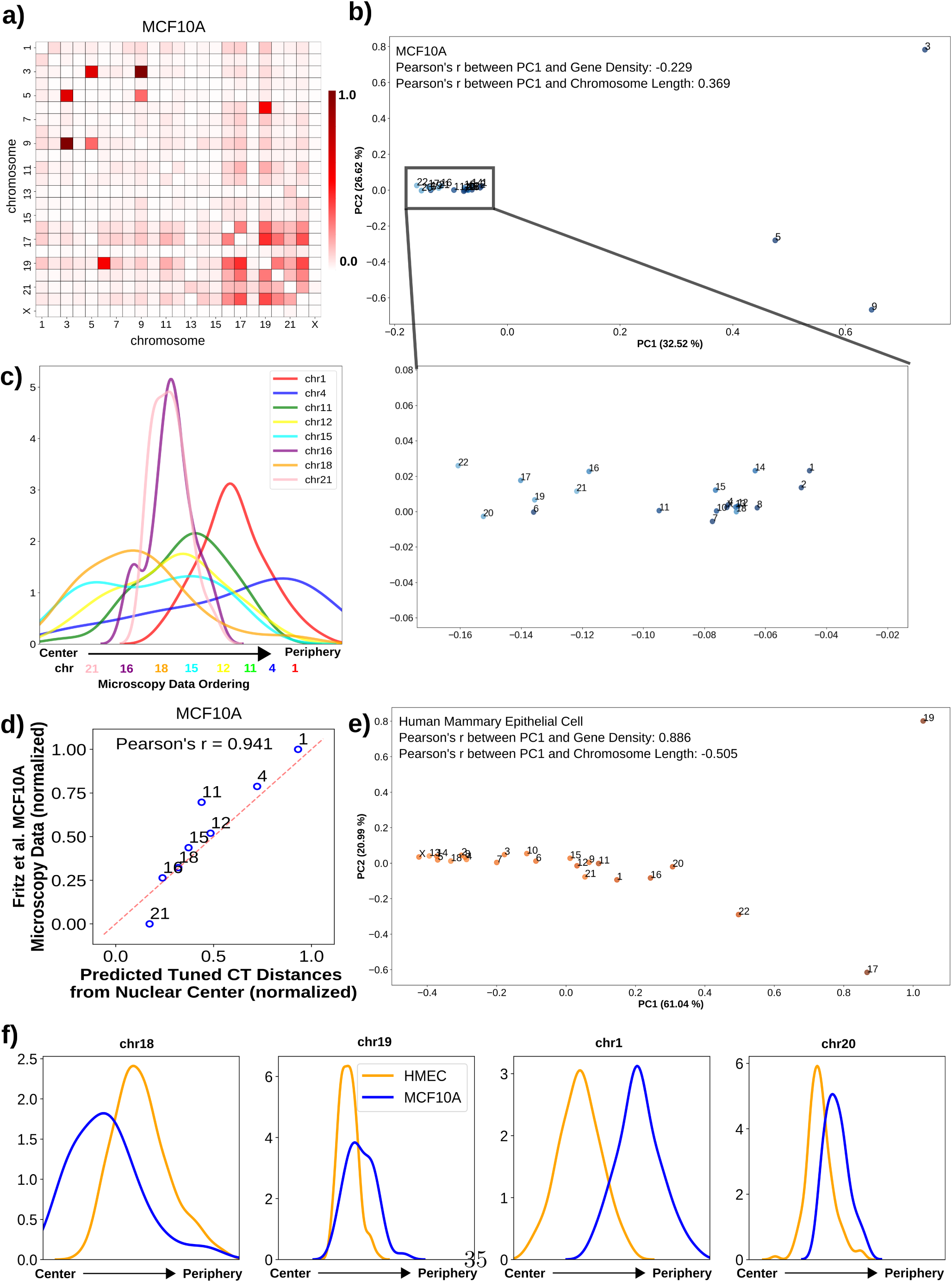
Epithelial cells show different radial CT organization in a nuclear morphology dependent manner. **a)** Pairwise inter-chromosomal strong interaction pattern matrix for MCF10A Hi-C data. **b, e)** 2D PCA projection of the pairwise inter-chromosomal strong interaction pattern matrices obtained from MCF10A (**b**) and HMEC (**e**) Hi-C data respectively. **c)** The radial distance profiles of 8 CTs (chr1, chr4, chr11, chr12, chr15, chr16, chr18 and chr21) for MCF10A obtained from the selected model cluster (# of models in the selected cluster: 48). Relative mean chromosome positions measured by microscopy shown below graph. **d)** Correlation between predicted tuned CT distance from Hi-C data and microscopy imaging data for MCF10A cells. **f)** The radial distance profiles of 4 CTs (chr18, chr19, chr1 and chr20) for both MCF10A and HMEC obtained from the respective selected model clusters (# of models in the selected clusters: for HMEC: 165).

Since epithelial cells can take different nuclear shapes depending on their properties in culture, we next applied our analysis to Hi-C data from another human normal mammary epithelial cell (HMEC) [70]. In this cell type, the PCA analysis of the pairwise inter-chromosomal strong interaction pattern inferred a gene density based distribution as represented in Fig. 5e. We hypothesize that this is related to the fact that these cells are closer to normal epithelium than MCF10A, which harbor chromosomal translocations, and that classical epithelial patterns would lead to a more spherical nucleus shape even in 2D culture, rather the flat, spreading growth pattern of MCF10A cells [72]. When the network modeling inferred radial distance profiles of four CTs - CT1, CT18, CT19 and CT20 were compared between MCF10A and HMEC (Fig. 5f), CT19 and CT20 showed density peaks at a similar radial position in both cell types. On the other hand, CT18 showed a preference for interior radial positions in MCF10A (length based) and for peripheral positions in HMEC (gene denisty based) and in case of CT1 that trend was the opposite. These results suggest that similar cell types can show different CT organization that correlates with their cell morphology, and Hi-C contact patterns can capture these differences.

### Variation between the radial arrangement of lymphoblastoid cells and neutrophils

To test the utility of our analyses on a cell having an irregular nuclear shape, we focused on neutrophils and obtained the corresponding Hi-C data from Javierre et al. [73]. Neutrophils have a multi-lobed nucleus with a toroid shaped genome where lobes are connected by thin filaments [74]. When we analyzed the pairwise inter-chromosomal strong interaction pattern matrix using PCA, we found that the PC1 values have a higher absolute correlation with gene density (Fig. 6b), though the effect is much weaker compared to GM12878 case, which is consistent with the divergence of a neutrophil from its precursor cell’s round nucleus shape. This is another indication that the Hi-C contact data provides more information about chromosome positioning than just the underlying length based or gene density based distribution. Based on microscopy imaging data, it has been reported that chr2 and chr18, which have drastic differences in their lengths, both occupy a position near nuclear envelope in neutrophil cells [75]. Upon inspecting the positions of those chromosomes along the PC1 axis in Fig. 6b, we found the two chromosomes in very close proximity near the left extreme of the PC1 axis. Although projection data does not infer the directionality of the ordering, based on its higher similarity with gene density, we can assume the left extreme as the periphery which contains mostly gene poor chromosomes. Furthermore, for those two chromosomes, we also observed similar trend in the network model-derived radial CT distance profiles as represented in Fig. 6c. The network model predicts a highly variable positioning of chr2, likely related to the overall dearth of strong contacts between chr2 and other chromosomes in the initial thresholded contact map. We observe that the predicted tuned CT positions for neutrophils are only weakly correlated with lymphoblastoid imaging data (Fig. 6d), revealing the ability of the model to predict different chromosome positions from different initial Hi-C contact data.

**Figure 6.**
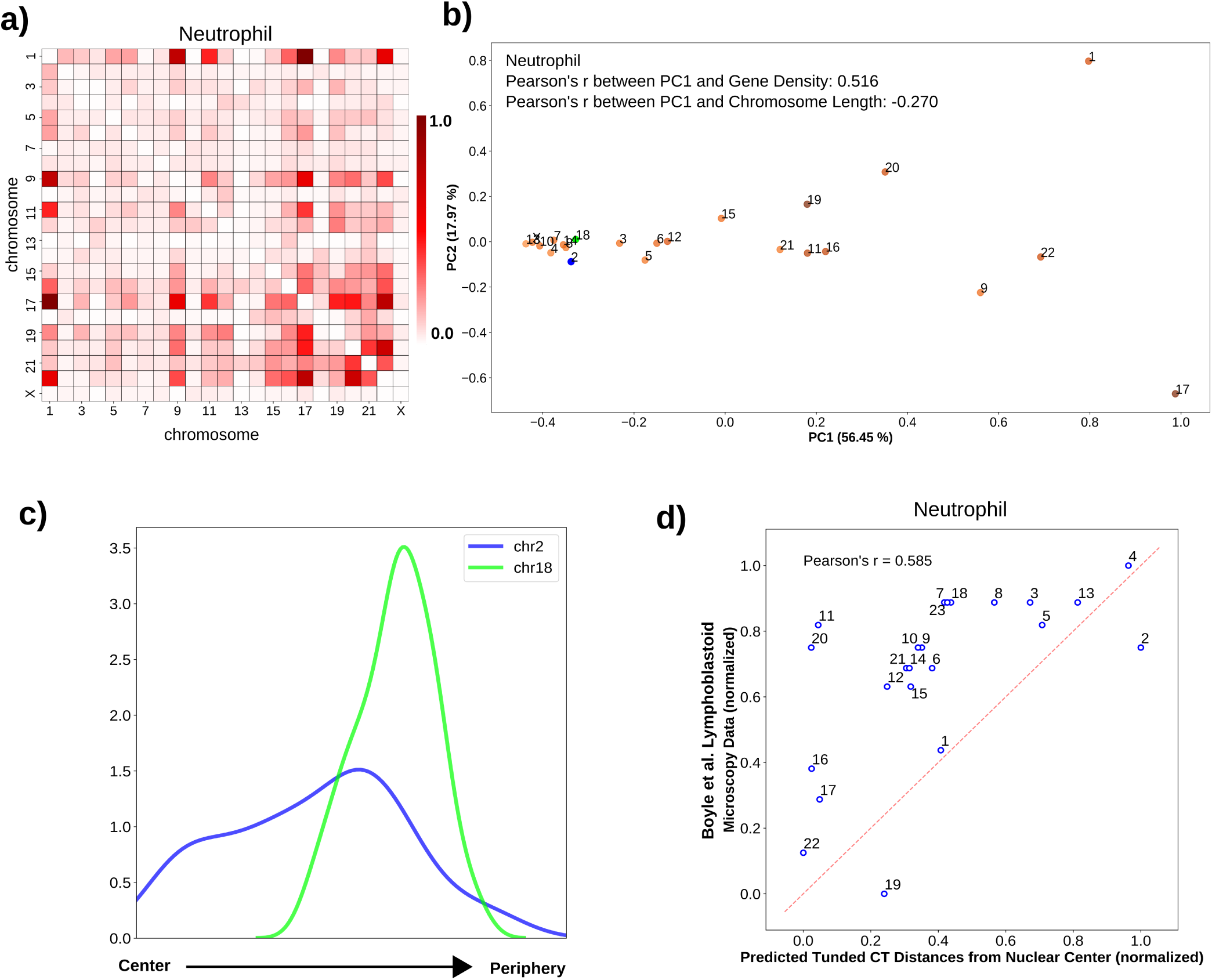
Radial arrangement of CTs in irregularly shaped neutrophil nuclei. **a)** Pairwise inter-chromosomal strong interaction pattern matrix for neutrophil Hi-C data. **b)** 2D PCA projection of the pairwise inter-chromosomal strong interaction pattern matrix obtained from neutrophil Hi-C data. In neutrophil cells, chr2 (blue color) and chr18 (green color) occupy radial positions near nuclear envelope. **c)** The radial distance profiles of 2 CTs (chr2, and chr18) in neutrophils obtained from the selected model cluster (# of models in the selected clusters: 100). **d)** Comparison of predicted tuned CT distances from neutrophil Hi-C data and lymphoblastoid microscopy imaging data shows important differences between these cell types.

## Discussion

In this study, our goal has not been only to predict one final set of CT positions from Hi-C data. Instead, we have explored what aspects of 3D chromosome radial positioning can and cannot be inferred with a series of direct and non-computationally intensive Hi-C contact analysis approaches that have not been previously used for this purpose.

Our results demonstrate that a straightforward statistical calculation (PCA) on the pattern of strong inter-chromosomal contacts can capture important biological features of CT radial positioning. With this approach, we not only can capture important patterns of gene density or chromosome length based ordering of chromosomes previously observed for very different cell types, but also can detect meaningful shifts of individual chromosomes in related cell types. We were able to infer changes in CT ordering in premature aging and senescence directly from contact data, and these changes are supported by previous microscopy results. We also note that this PCA ordering approach is actually quite robust to differences in Hi-C data quality and depth. While our subsequent network graph layout approach was somewhat sensitive to different quality metrics of different Hi-C replicates, the PCA ordering of chromosomes was robust to different levels of noise in these datasets.

We have also demonstrated that a network graph layout algorithm approach can generate an ensemble of models that capture the experimentally validated mean and variation of CT positions in a cell type. We demonstrate that these approaches can be applied to a variety of cell types and can even detect differences in underlying CT radial distributions between highly related cell types (two different breast epithelial cell types). Interestingly, for these cell types, the difference in CT distribution corresponds to documented differences in nucleus geometry of these cell types when grown in culture. This suggests that not only do spherical and elliptical nuclei exhibit different CT organization in completely different cell lineages (lymphoblast vs. fibroblast), as previously documented, but that even cells within the same overall type may have different CT positioning associated with their nucleus shape. This result emphasizes the importance of approaches to detect changes in CT ordering directly from a given cell type in its particular condition, rather than assuming that a measurement in one circumstance will define CT organization principles across, for example, all epithelial cells. Our rapid approaches to inferring CT distribution makes screening across such varieties of datasets more feasible, as a complement to more intensive computational models or microscopic measurements.

In addition to the validation of these approaches that makes them useful for future applications, our results also show that while Hi-C contacts are useful for inferring a relative ordering of radial chromosome positions, inferring absolute ordering often requires an external reference point. We observe this in that the PCA based ordering of chromosomes can represent either direction (interior-exterior or vice versa) and that some clusters of network models display the same chromosome relative ordering in reverse orientation. This is an important factor to consider in any model that attempts to use Hi-C contacts in isolation to generate 3D structure models.

The analysis approach used in this paper is highly flexible due to its modular nature and can be integrated with different kinds of genome analysis applications. For example, both PCA ordering and inferred tuned positions can be used to characterize the changes in the radial CT organization in a perturbed (e.g. relocation of CTs during DNA damage response [76]) or diseased cell (e.g. mislocalization of CT18 and CT19 in lamin B2 depleted colorectal cancer cells [77]) compared to a control cell, which in turn allows us to study how these changes affect higher-order chromatin structure. The method also generates a population of 3D network structures, which can further be used to characterize inter-chromosomal dynamics and can be compared to single-cell Hi-C results.

## Methods

The schematic representation of our whole analysis approach is given in Fig. 7.

**Figure 7.**
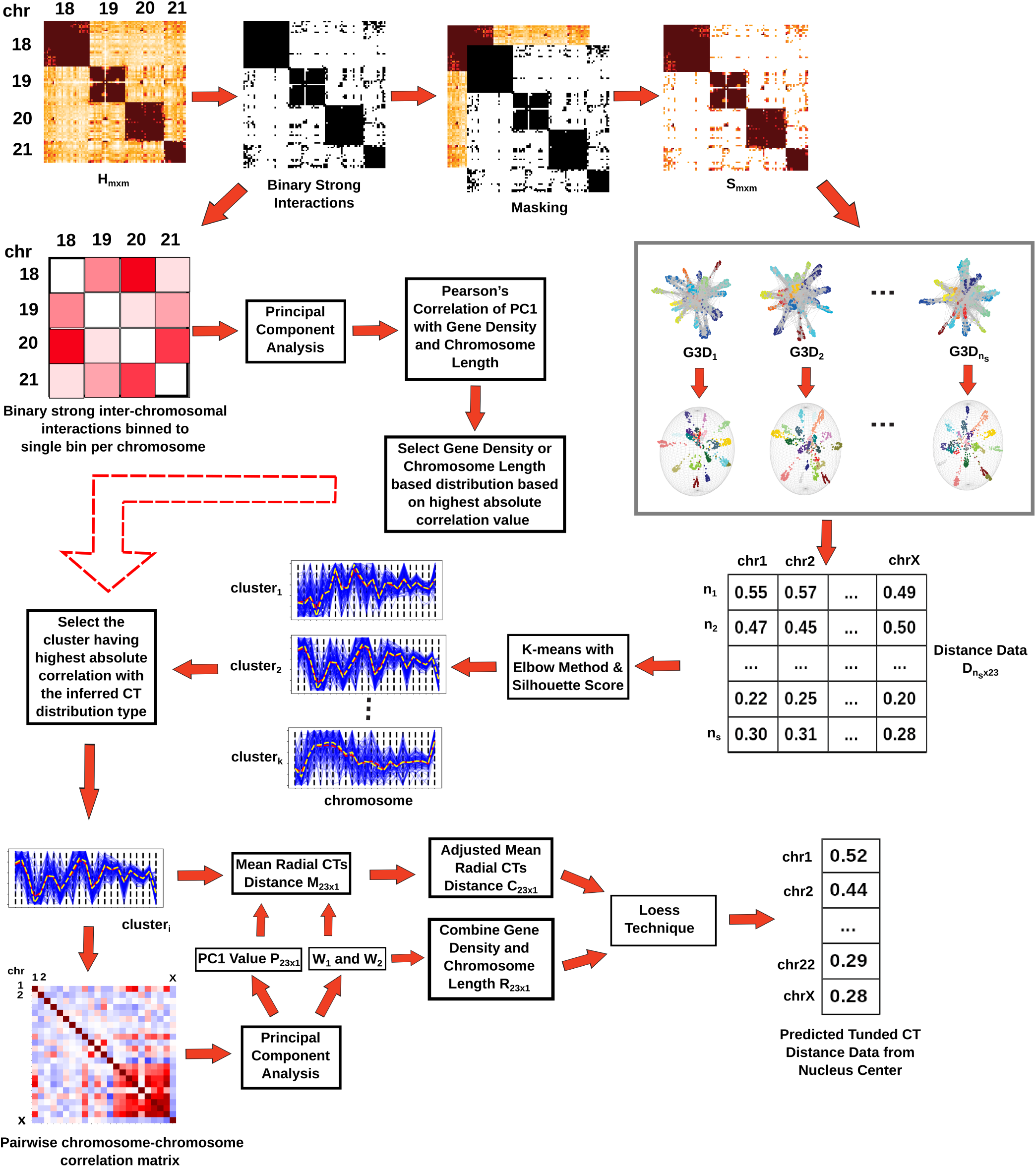
Overview of the computational analysis approaches. At the first step, the whole genome Hi-C matrix is thresholded in order to extract strong interactions. Next, the pairwise inter-chromosomal strong interaction pattern matrix obtained from the thresholded Hi-C data is analyzed using PCA to determine the underlying CT distribution type. The radial CT organization is also modeled from the thresholded interactions by generating multiple 3D network model configurations. Out of those configurations, clusters of models having similar radial arrangements are identified and the particular cluster matching the inferred CT distribution type from PCA is selected for further downstream analysis. Finally, the network modeling inferred radial organization can be tuned by incorporating the chromosome gene density and length properties.

### Pre-processing of Hi-C data

All original Hi-C data sources for this study are listed in Supplementary Table 1. We mapped the data to hg19 to obtain a set of unique valid interacting fragment pairs and then binned the data at a resolution of 2.5 Mb, where each bin of the contact matrix represents the interaction frequency between a pair of 2.5 Mb genomic regions. We then perform two pre-processing steps on the Hi-C matrix. First, to remove biases related to GC content and cut site frequency, the raw contact matrix is normalized using the ICE technique [78]. Next, the unmappable (repetitive) genomic regions are removed from the normalized Hi-C matrix. Also, we do not consider chrY in our analysis since this chromosome is not present in both male and female cell types, and we remove those corresponding bins from the Hi-C matrix. This modified Hi-C matrix is used for further downstream processing.

Let *H*_*m×m*_ represent a square symmetric Hi-C contact matrix where each row and column correspond to genomic regions (bins) of specific size, with each element *h*_*ij*_ representing the normalized number of contacts between *i*^*th*^ and *j*^*th*^ bins.

### Determining a threshold to capture strong interactions in Hi-C matrix

After prepossessing the Hi-C contact data, we proceed to identify the strong interactions which have the best potential to infer cell-type specific radial organization of the CTs, distinct from the levels of average background noise. To do this, we define a cutoff limit and apply that to *H*_*m×m*_, to obtain a matrix of only filtered strong interactions *S*_*m×m*_.

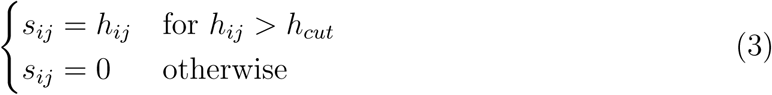

where *s*_*ij*_ is the element of matrix *S*_*m×m*_, which represents the normalized number of contacts between *i*^*th*^ and *j*^*th*^ bins and *h*_*cut*_ is the cutoff limit. For 2.5 Mb genomic resolution, the value of cutoff limit is set to: *h*_*cut*_ = 95^*th*^ percentile of all the genome-wide interactions from *H*_*m×m*_. The significance of and reason for choosing this 95^*th*^ percentile cutoff is discussed in the **Results** section. Due to the application of this cutoff limit *h*_*cut*_, the resultant matrix *S*_*m×m*_ contains mostly the intra-chromosomal interactions and a few inter-chromosomal interactions. In addition, to detect significant chromosomal interactions, we apply FitHiC tool with default parameters to the genome-wide Hi-C contact data, binned at 2.5 Mb resolution. From those significant intra- and inter-chromosomal interactions detected, we further select highly significant interactions for analysis by applying a q-value cutoff - 10^−2^ (for intra-chromosomal) and 10^−12^ (for inter-chromosomal). The reason behind selecting a very stringent q-value cutoff for inter-chromosomal interactions is to make the number of significant inter-chromosomal interactions from FitHiC comparable to the strong interactions from our method.

### Simulating random ligation Hi-C from original Hi-C data

Random ligation Hi-C data are simulated by taking an original raw/non-normalized Hi-C contact map, binned at 2.5 Mb, and shuffling the bins (including the diagonal) of this matrix five times. Then, the shuffled matrix is passed through the ICE normalization step. The reason for generating the random ligation matrix from the original matrix by random shuffling is to ensure that both the contact matrices have an equal number of total reads and a similar dynamic range of values, ensuring a matched comparison of the results from real and random Hi-C data.

### Identifying CT radial distribution patterns with PCA transformation on the pairwise strong inter-chromosomal interaction pattern matrix

For our most direct approach to infer a radial organization pattern from Hi-C data, we begin by creating a pairwise strong inter-chromosomal interaction pattern matrix. For the chosen *h*_*cut*_ value, the number of bin pairs passing the threshold are summed between each pair of chromosomes to a single bin (Eqn. 4). This sum thus includes a count of the number of bin pairs that passed a threshold, rather than the total number of interactions in each included bin.

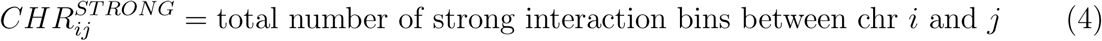

Then, we exclude cis contacts and look only at the inter-chromosomal component of this pairwise strong interaction pattern (Eqn. 5).

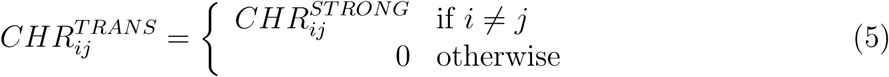

To capture the major interaction trends from this matrix, we apply principal component analysis (PCA) to this pairwise strong inter-chromosomal interaction pattern. We then calculate the projection of all chromosomes onto PC1, and we find that the ordering of chromosomes along this PC can capture the radial organization ordering of the chromosomes.

### Constructing 3D network graphs from Hi-C matrix

The thresholded strong interaction matrix *S*_*m×m*_ is treated as a weighted adjacency matrix to generate an undirected weighted graph *G*, where nodes represent genomic bins and the number of interactions between each pair of bins is used as the edge weight. Next, we apply a 3D Fruchterman-Reingold (FR) force based layout to draw the undirected Hi-C graph in 3D space. This graph drawing layout adds an attractive force between the connected nodes and creates repulsion between the nodes that are not connected. Along with this, gravitational force is used in this layout to pull the nodes towards the center. The rationale behind using the FR layout is that it uncovers the intrinsic structure of the network, as the strong interactions only matrix *S*_*m×m*_ has a large number of intra-chromosomal interactions and fewer inter-chromosomal interactions. The resulting 3D network graph infers the radial organization of the CTs inside the nucleus. Next, *n*_*s*_ number of 3D network graphs - 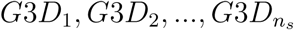 are generated by performing independent runs of the FR algorithm with different random initial configurations to model the variability in the radial CT organization.

### Fitting a geometrical structure to the 3D network graphs

For each 3D network graph, the next objective is to find the distance of each CT from the center of that network graph. This step is performed in two parts.

In the first part, given a 3D network graph, the 3D Cartesian coordinates of the nodes are extracted and then Khachiyan’s algorithm is used to find a minimum volume ellipsoid enclosing the set of nodes [79]. The center of the fitted object is calculated and assigned as the nucleus center. Along with this, the center of mass of each of the individual CTs is calculated from the coordinates of the nodes of the network graph.

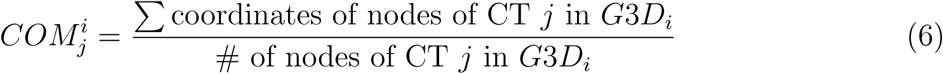

where 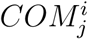 represents the center of mass of a particular CT *j* in the structure *G*3*D*_*i*_ and *j* ∈ {1, 2, …, 22, *X*}.

After obtaining the center of mass of each CT and nucleus center, the Euclidean distance is calculated between the center of mass of each CT and the nucleus center. Also, to remove the heterogeneity that arises from different minimum volume ellipsoid fits of different 3D network structures, for each structure the distance values are normalized in the [0,1] range using Min-max normalization. In this way, by iterating over all of the 3D network graphs, for each CT, *n*_*s*_ distances from the center of the nucleus are obtained. This distance information is represented in a matrix 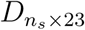 of size *n*_*s*_ × 23, where rows represent *n*_*s*_ 3D network graphs and columns correspond to 23 CTs (22 autosomes and one X chromosome), and each matrix entry *d*_*ij*_ is the distance of the CT *j* from the nucleus center of network graph *G*3*D*_*i*_.

The parameter *n*_*s*_ represents the number of configurations required to mimic the heterogeneity in CT arrangement originating during cell division within the same cell type. To estimate this parameter, for different values of *n*_*s*_ we compare the fitted distance profiles of all CTs between GM12878 and BJ1-hTERT cells using non-parametric hypothesis testing - a Two-sided Mann-Whitney U test. From the statistical test results, it can be observed that with increasing *n*_*s*_ value, more chromosomes show significant differences in their CT distance profiles between lymphoblastoid and fibroblast cells, reaching a maximum at *n*_*s*_ = 1000 (Supplementary Fig. 9). Hence, we set the *n*_*s*_ parameter to 1000 in our analysis.

### Identifying the specific cluster of network models having meaningful CT organization

The network modeling approach produces 3D graph models with heterogeneous CT organization, as intended, but we find that some groups of models may capture an inverted ordering of some groups of CTs or all CTs (reversing central to peripheral distances). Thus, rather than blindly averaging all models together, we first identify these different clusters of organization patterns and then choose for further consideration the cluster of models that follows the radial CT organization distribution captured by the PCA analysis described above. We perform K-means clustering [80] with deterministic initialization on the distance matrix 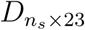 by treating the models (rows) as samples and chromosomes (columns) as features. The initial centroid positions for the clustering technique are calculated using the algorithm from Nazeer et al. [81]. In addition to that, to estimate optimal number clusters for the K-means, we use a modified elbow method approach. Here, first we calculate inertia [82] which represents within cluster sum of squares for different increasing number of clusters and detect the elbow of the inertia curve with the help of the algorithm from Satopaa et al. [83]. After detecting the elbow, again we calculate the average silhouette score [82] for each of the different number of clusters and select the point as optimal number of clusters which will have the highest average silhouette score in the vicinity of the elbow point (two points upstream of elbow, two points downstream and the elbow itself). As our set of predictions for further analysis, we take the cluster whose mean radial CT positions shows the highest absolute correlation with the CT distribution type obtained by the PCA transformation of the pairwise inter-chromosomal strong interaction pattern matrix.

### Gene density and chromosome length based tuning of consensus radial organization of CTs

The last aspect of our approach considers how the predicted averaged radial CT distances can be further tuned using two chromosomal properties - gene density and chromosome length. Human chromosome gene density (genes/ sequenced Mb) is obtained from “Short guide to the human genome” [84] and length information from UCSC Genome Browser hg19 [85]. But before discussing the steps involved with this tuning procedure, let us assume, 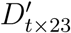 represents the selected model cluster based on inferred CT distribution type. This cluster contains *t* number of network models and the average CT distance of each chromosome across all of the *t* models is denoted by the column vector *M*_23*×*1_. After having the selected model cluster 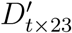 we calculate a pairwise chromosome correlation matrix of size 23×23 from that and perform PCA transformation on that correlation matrix. The PC1 value obtained from this transformation represents the major separation of the chromosomes based on pairwise interactions and is denoted by the column vector *P*_23*×*1_. Once we have the two column vectors -*M*_23*×*1_ and *P*_23*×*1_ representing the average radial position of the CTs and their separation respectively, we combine them using a weighted averaging technique as per Eqn. 7.

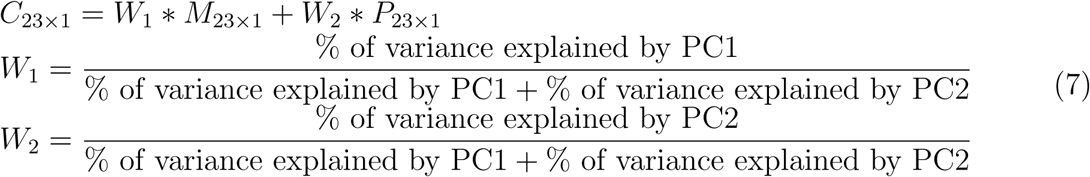

where *C*_23*×*1_ represents the resultant combined column vector and *W*_1_ and *W*_2_ denote the weights calculated from the percentage of the the variance explained by PC1 and PC2.

Next, in order to model the effect of both chromosome length and gene density, we combine these two properties using weighted averaging as described in the Eqn. 8. The first and second components on the right hand side of each of the equations are related to the normalized gene density and normalized chromosome length respectively. Also, we use the same set of weights *W*_1_ and *W*_2_ in this equation but the weights are ordered based on the inferred CT distribution type. For example, if the inferred CT distribution follows a gene density based radial positioning, in that case the component having gene density will have the higher weight *W*_1_ and vice verse

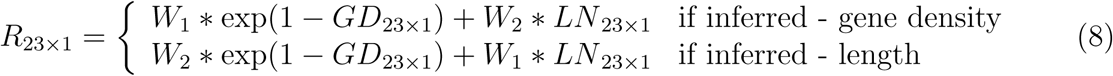

Here, *GD* and *LN* are column vectors of size 23 ×1 and contain the gene density and length of each chromosomes respectively.

Following the chromosomal properties modeling procedure, in the final step of the tuning, the modified consensus radial distance of the CTs *C*_23*×*1_ is locally smoothed based on *R*_23*×*1_ using the Loess technique. The radial distance of the CTs obtained from this step represents the tuned arrangements of the CTs inside the nucleus.

## Supplementary Materials for

**Supplementary Table 1:** Dataset description

| Cell Line | Type | Nuclear Shape | Hi-C Data | GEO or EGA Accession Number | Microscopy Data |
| --- | --- | --- | --- | --- | --- |
| GM12878 | Lymphoblastoid | Spherical | Sanders et al. (2019) | GSE136899 | Boyle et al. (2001) |
| GM12878 Rao | Lymphoblastoid | Spherical | Rao et al. (2014) | GSE63525 | Boyle et al. (2001) |
| BJ1-hTERT | Skin Fibroblast | Ellipsodial | Sanders et al. (2019) | GSE136899 | Bolzer et al. (2005) |
| BJ-5ta | Skin Fibroblast | Ellipsoidal | Sanders et al. (2019) | GSE136899 | Bolzer et al. (2005) |
| MCF10A | Breast Epithelium | Ellipsoidal | Barutcu et al. (2015) | GSE66733 | Fritz et al. (2014) |
| HMEC | Breast Epithelium | Spherical | Rao et al. (2014) | GSE63525 | - |
| Neutrophil | Neutrophil | Multi Lobed | Javierre et al. (2016) | EGAS00001000327 | - |
| WI38-hTERT | Lung Fibroblast | Ellipsoidal | Chandra et al. (2015) | ENA PRJEB8073 | - |
| WI38-hTERT Senescent | Lung Fibroblast | Ellipsoidal | Chandra et al. (2015) | ENA PRJEB8073 | - |
| HGPS Patient p19 | Lymphoblastoid | Ellipsoidal | McCord et al. (2013) | GSE41763 | - |

**Supplementary Figure 1.**
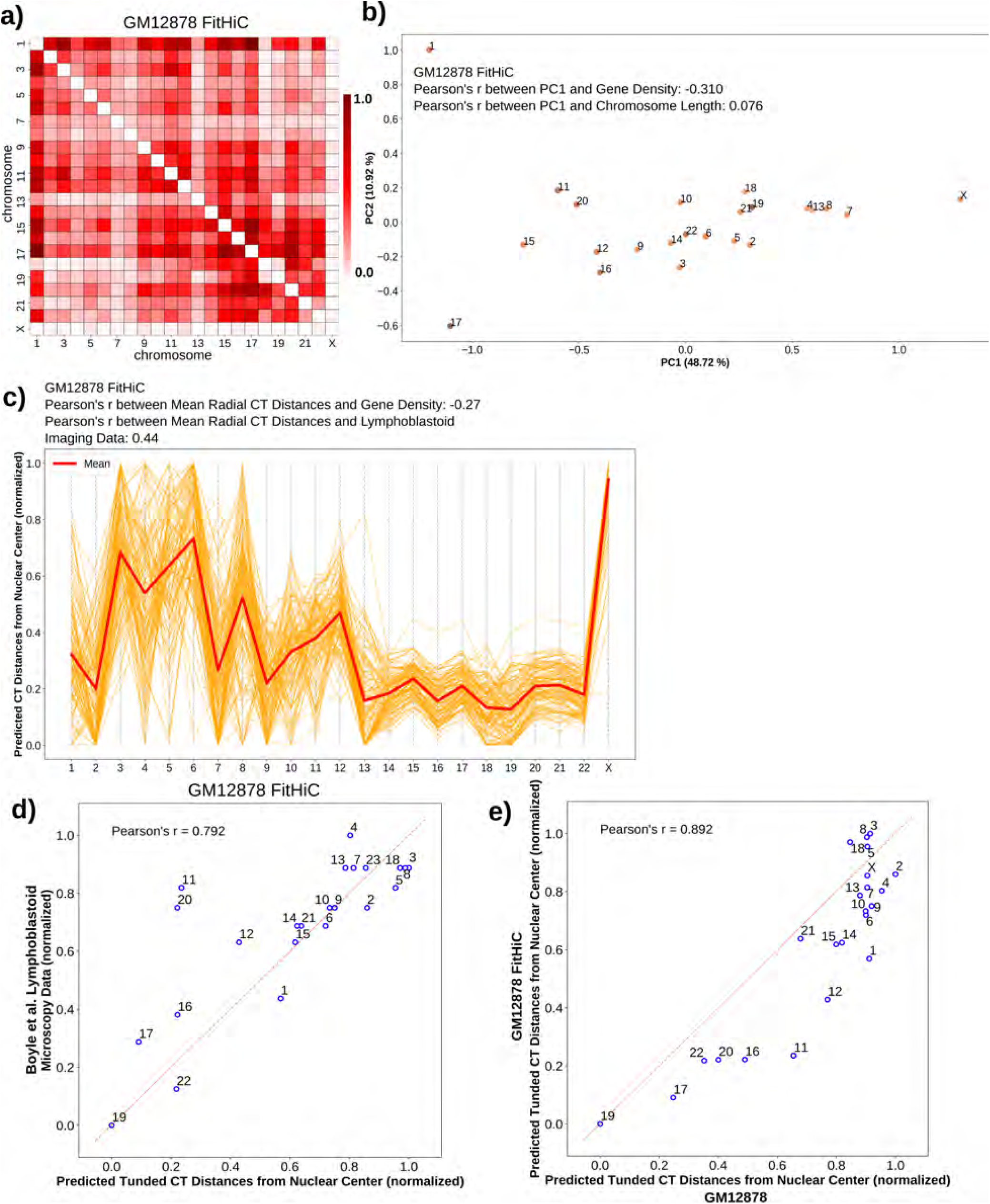
Radial CT arrangement of GM12878 (FitHiC). **a)** Pairwise inter-chromosomal significant interaction pattern matrix derived from FitHiC for GM12878 Hi-C data. **b)** 2D PCA projection of the FitHiC pairwise inter-chromosomal significant interaction pattern matrix obtained from GM12878 Hi-C data. **c)** Network modeling generated model cluster for FitHiC GM12878 contacts, selected based on inferred CT distribution type. **d)** Correlation between predicted tuned CT distance from GM12878 (FitHiC) and lymphoblastoid microscopy imaging data. **f)** Correlation between predicted tuned CT distances obtained from GM12878 (FitHiC) and GM12878 (strong interactions)

**Supplementary Figure 2.**
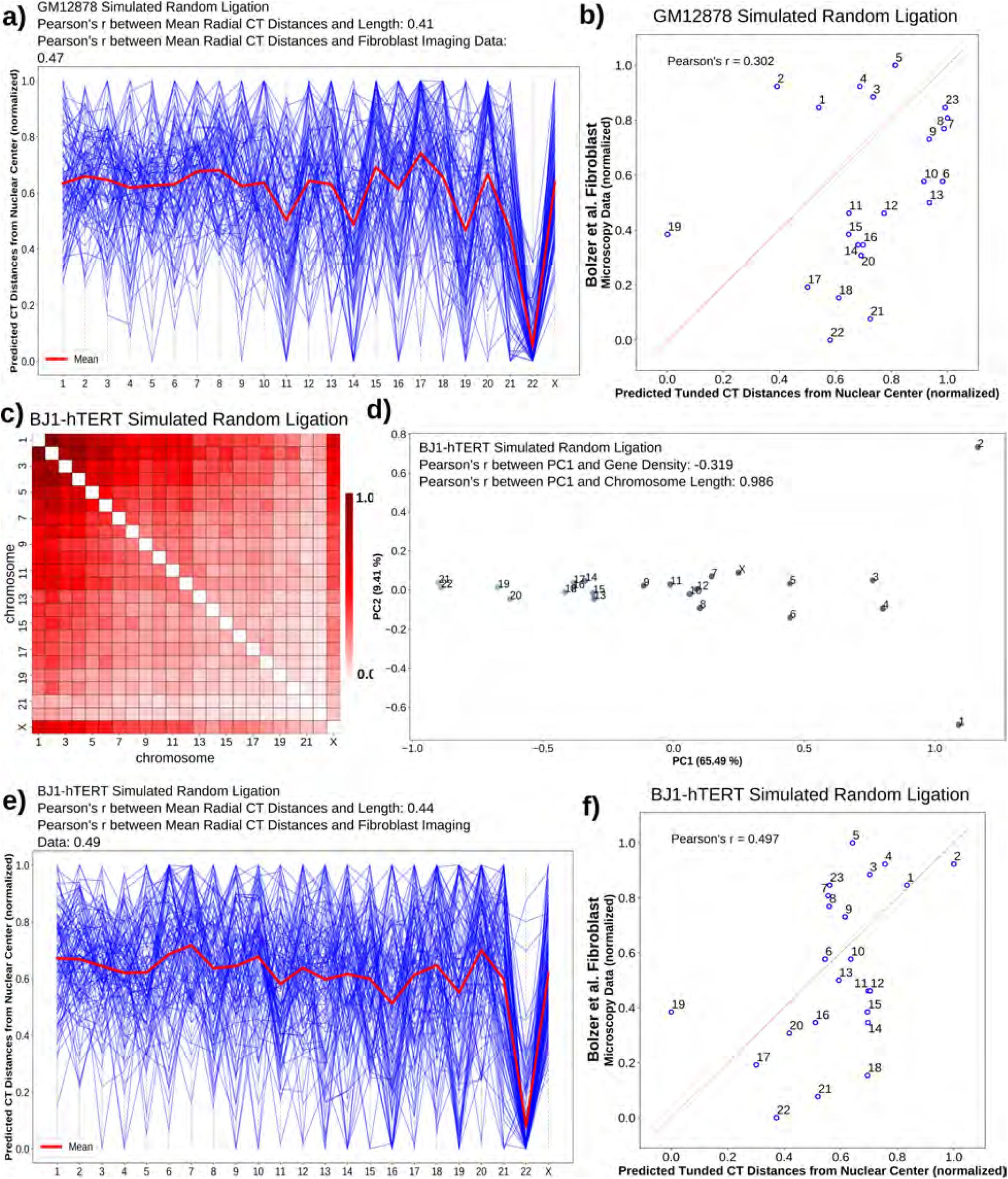
Radial CT arrangements inferred from GM12878 and BJ1-hTERT simulated random ligation data. **a)** Network modeling cluster for GM12878 simulated random ligation, selected based on inferred CT distribution type. **b)** Correlation between predicted tuned CT distance from GM12878 simulated random ligation Hi-C data and lymphoblastoid microscopy imaging data. **c)** Pairwise inter-chromosomal strong interaction pattern matrix for BJ1-hTERT simulated random ligation Hi-C data. **d)** 2D PCA projection of the pairwise inter-chromosomal strong interaction pattern matrix obtained from BJ1-hTERT simulated random ligation Hi-C data. **e)** Network modeling cluster for BJ1-hTERT simulated random ligation, selected based on inferred CT distribution type. **f)** Correlation between predicted tuned CT distance from BJ1-hTERT simulated random ligation Hi-C data and fibroblast microscopy imaging data.

**Supplementary Figure 3.**
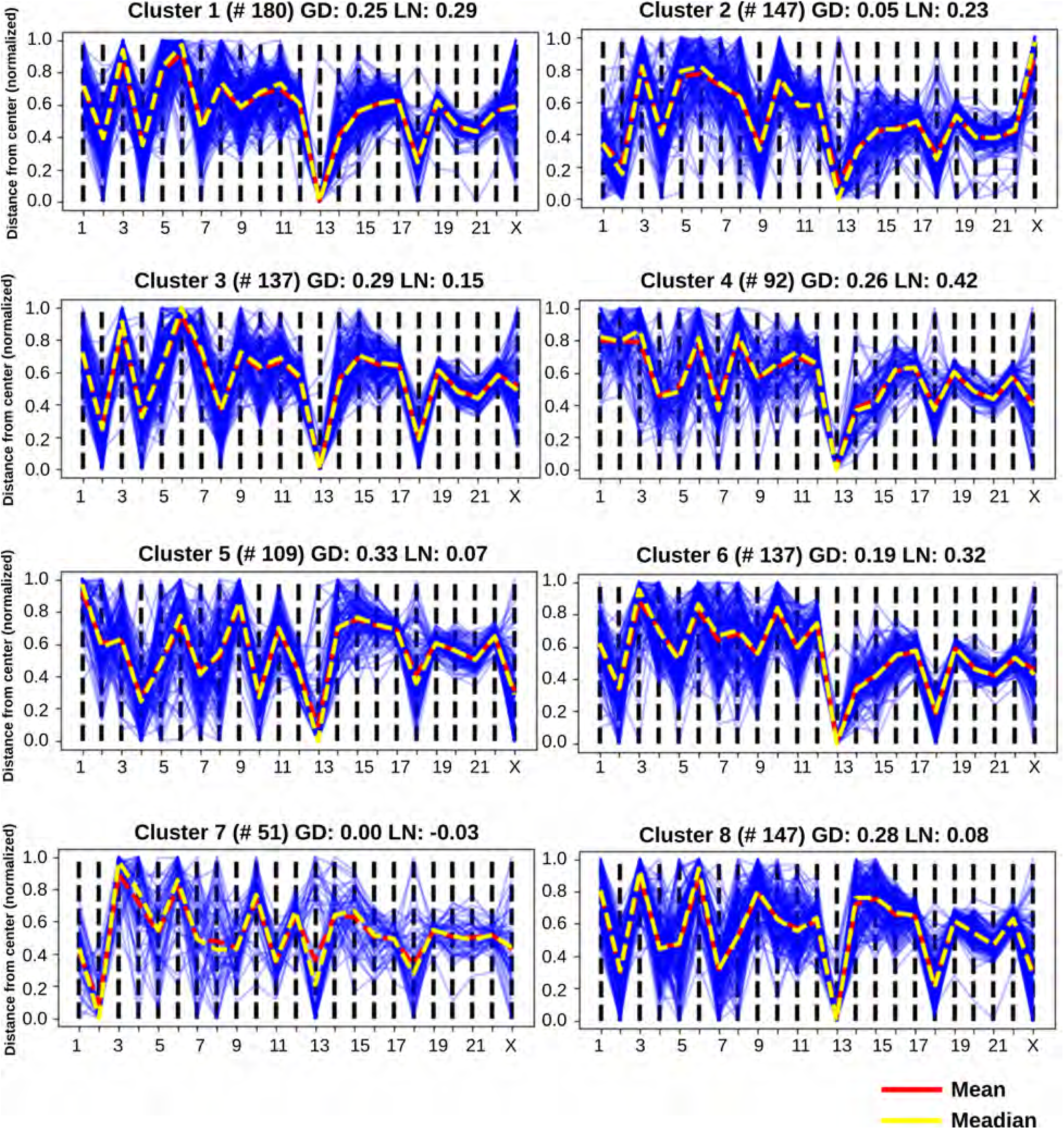
Network modeling generated clusters for GM12878. For each cluster, the Pearson’s correlation of mean radial CT distances with gene density (GD) and chromosome length (LN) are shown on the top of the cluster. Number of models in each cluster is indicated in parentheses above each cluster.

**Supplementary Figure 4.**
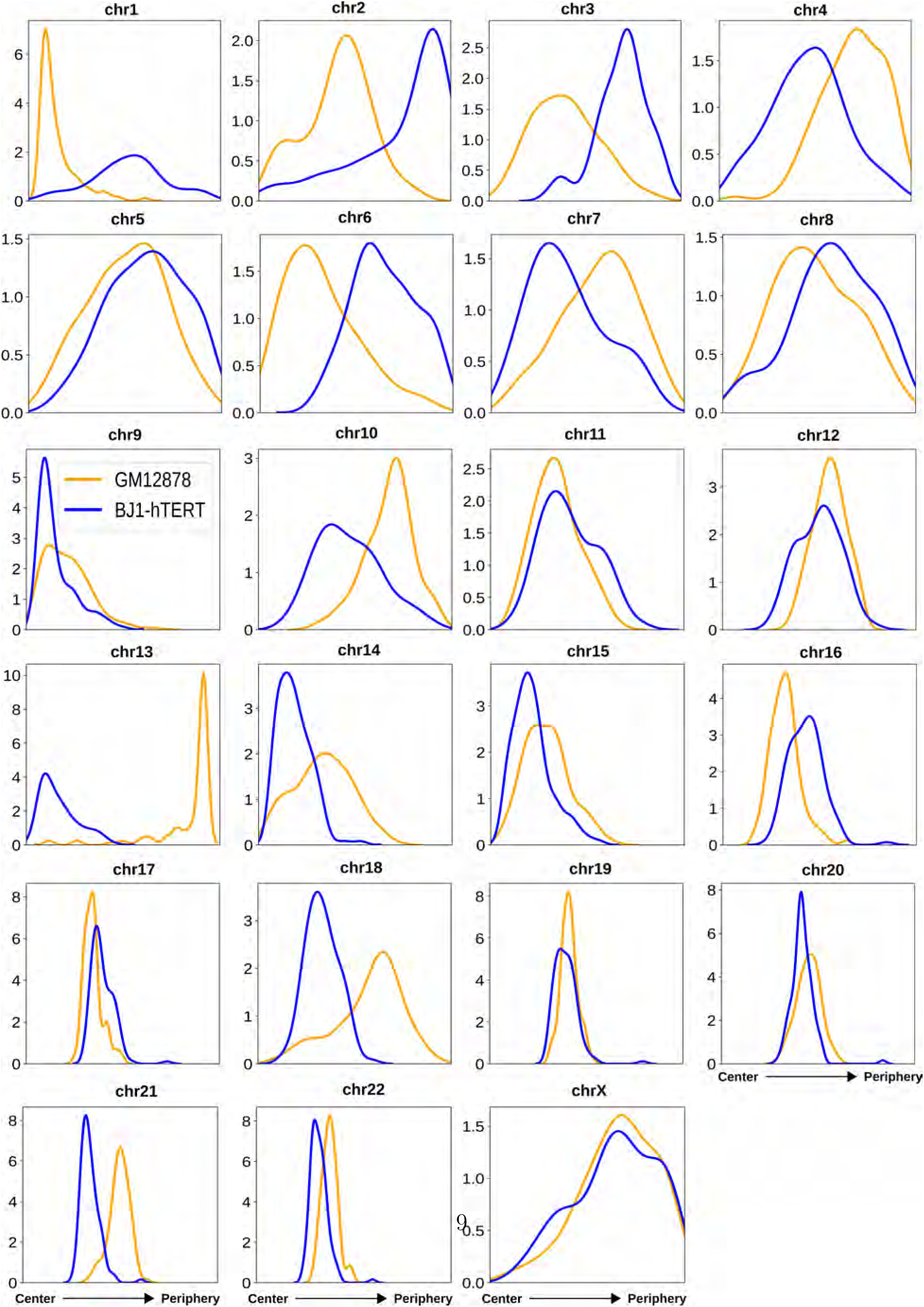
The radial distance distributions of 23 CTs for both GM12878 and BJ1-hTERT obtained from the respective selected model clusters based on inferred CT distribution types (# of models in the selected clusters-GM12878: 109, and BJ1-hTERT: 122). All distributions are ordered left to right from center to periphery.

**Supplementary Figure 5.**
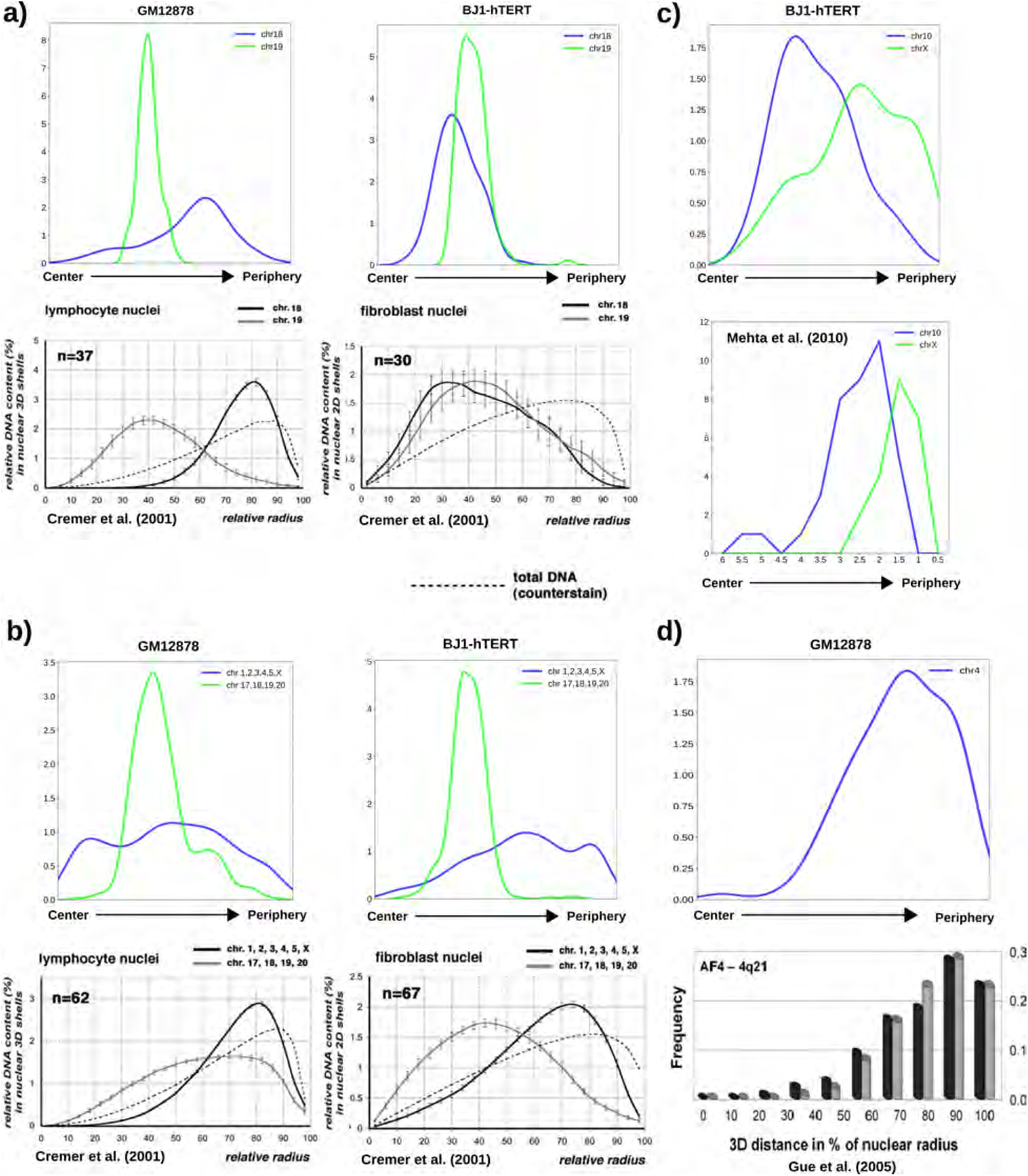
Comparison of network modeling generated radial CT distance profiles with experimentally obtained distributions. **a-b)** The radial distance profiles of CT18 and CT19 **(a top)** and short and long chromosomes **(b top)** for GM12878 and BJ1-hTERT obtained from the network model clusters in this current work. These distributions are similar to the radial arrangement previously measured by microscopy for CT18 and CT19 **(a bottom)** and short and long chromosomes **(b bottom)** in lymphocyte (2D arrangement) and fibroblast (3D arrangement) nuclei. Panel **(a-b)** bottom parts are adapted from Cremer et al. (2001). Copyright Springer Nature. Used with permission. **c) Top** - The radial distance profiles of CT10 and CTX for BJ1-hTERT obtained from our network model cluster. **Bottom** -The distance distribution of CT10 and CTX from the nuclear periphery in human dermal firbroblasts obtained using 3D FISH. Panel **(c)** bottom part is created with data obtained from Mehta et al. (2010). **d) Top** - The radial distance profiles of CT4 for GM12878 obtained from our network modeling cluster. **Bottom** - Radial distance profiles of AF4 gene, which is located on chr4, from the nucleus center in two lymphoblastic cell lines - NALM-6 (black) and IL-9 (grey). Panel **(d)** bottom part is adapted from Gué et al. (2005), copyright John Wiley and Sons, used with permission.

**Supplementary Figure 6.**
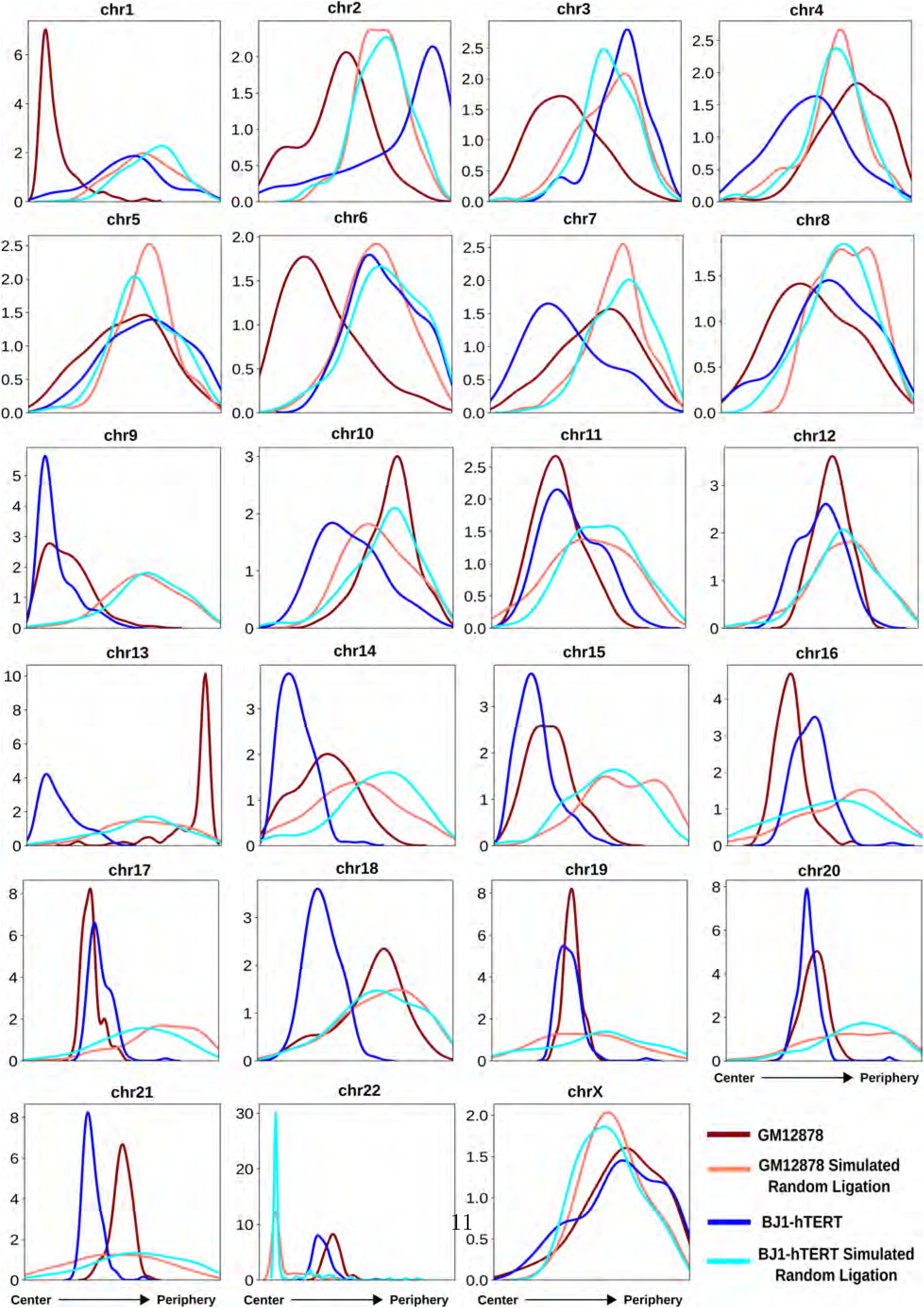
The radial distance distribution profiles of 23 CTs for GM12878, GM12878 simulated random ligation, BJ1-hTERT and BJ1-hTERT simulated random ligation obtained from the respective selected model clusters based on inferred CT distribution types (# of models in the selected clusters: for GM12878 - 109, GM12878 simulated random ligation - 95, BJ1-hTERT - 122, and BJ1-hTERT simulated random ligation - 113).

**Supplementary Figure 7.**
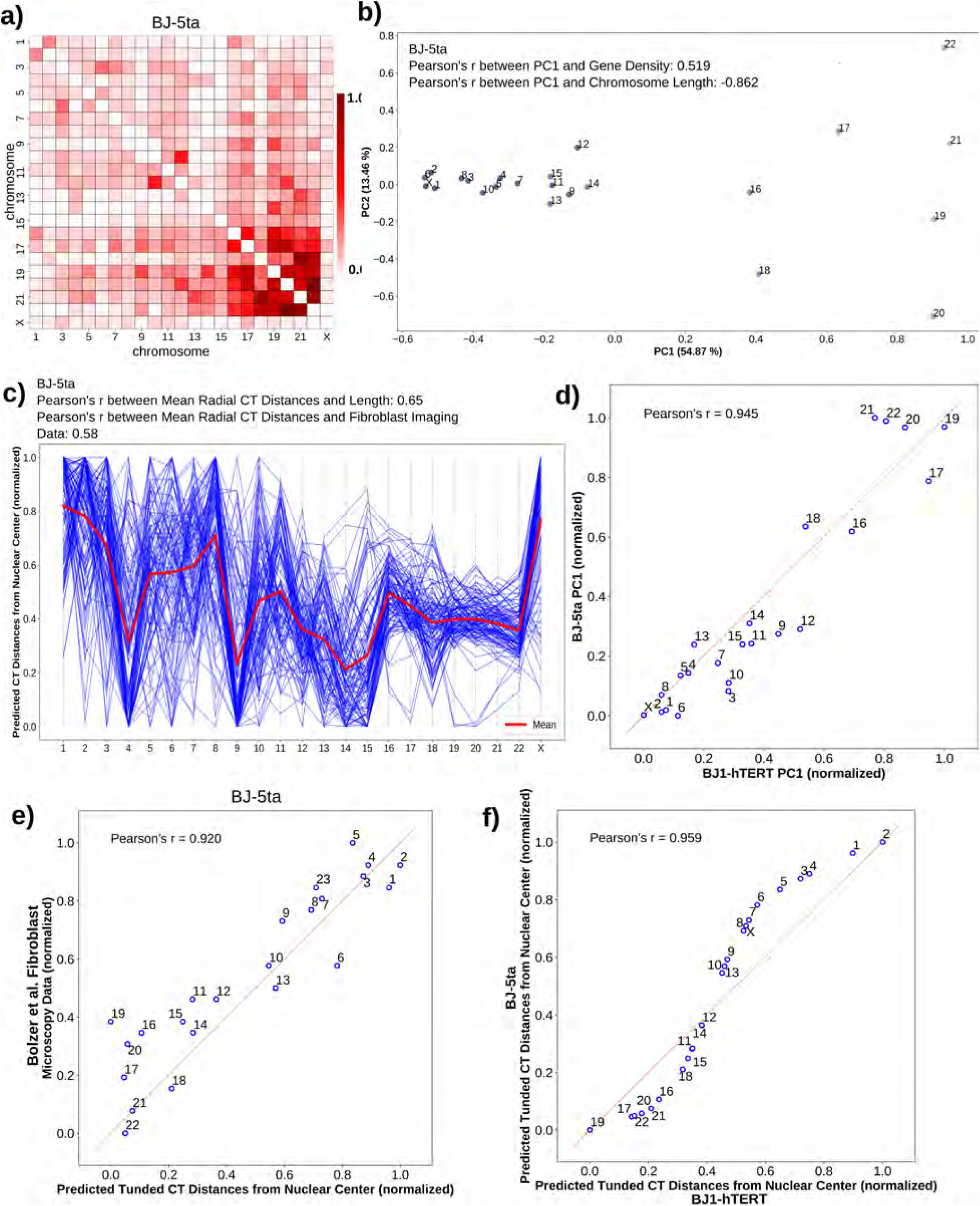
Radial arrangement of BJ1-hTERT and BJ-5ta follow similar CT distribution patterns. **a)** Pairwise inter-chromosomal strong interaction pattern matrix for BJ-5ta Hi-C data. **b)** 2D PCA projection of the pairwise inter-chromosomal strong interaction pattern matrix obtained from BJ-5ta Hi-C data. **c)** Network modeling generated model cluster for BJ-5ta, selected based on respective inferred CT distribution type (# of models in the selected clusters: 115). **d)** Correlation of BJ1-hTERT PC1 values with BJ-5ta PC1 values obtained from the PCA transformation of the respective pairwise inter-chromosomal strong interaction pattern matrices. **e)** Correlation between predicted tuned CT distance from BJ-5ta Hi-C data and fibroblast microscopy imaging data. **f)** Correlation between predicted tuned CT distances obtained from BJ1-hTERT and BJ-5ta.

**Supplementary Figure 8.**
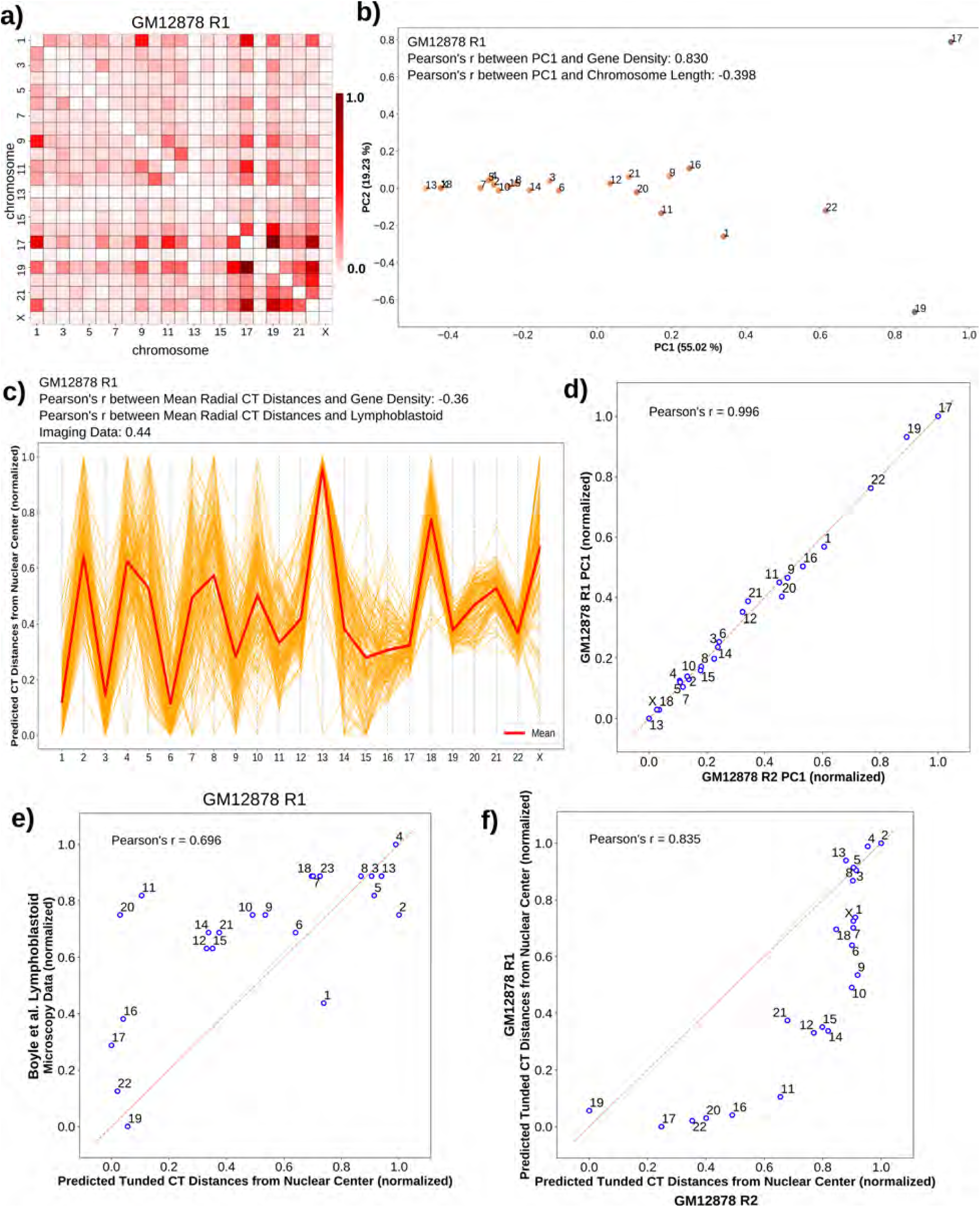
Radial arrangement of GM12878 replicates R1 and R2 show similar CT distribution pattern. **a)** Pairwise inter-chromosomal strong interaction pattern matrix for GM12878 R1 Hi-C data (R2 used in main figures). **b)** 2D PCA projection of the pairwise inter-chromosomal strong interaction pattern matrix obtained from GM12878 R1 Hi-C data. **c)** Network modeling generated model cluster for GM12878 R1, selected based on respective inferred CT distribution type (# of models in the selected clusters: 142). **d)** Correlation of GM12878 R2 PC1 values with GM12878 R1 PC1 values obtained from the PCA transformation of the respective pairwise inter-chromosomal strong interaction pattern matrices. **e)** Correlation between predicted tuned CT distance from GM12878 R1 Hi-C data and lymphoblastoid microscopy imaging data. **f)** Correlation between predicted tuned CT distances obtained from GM12878 R2 and GM12878 R1

**Supplementary Figure 9.**
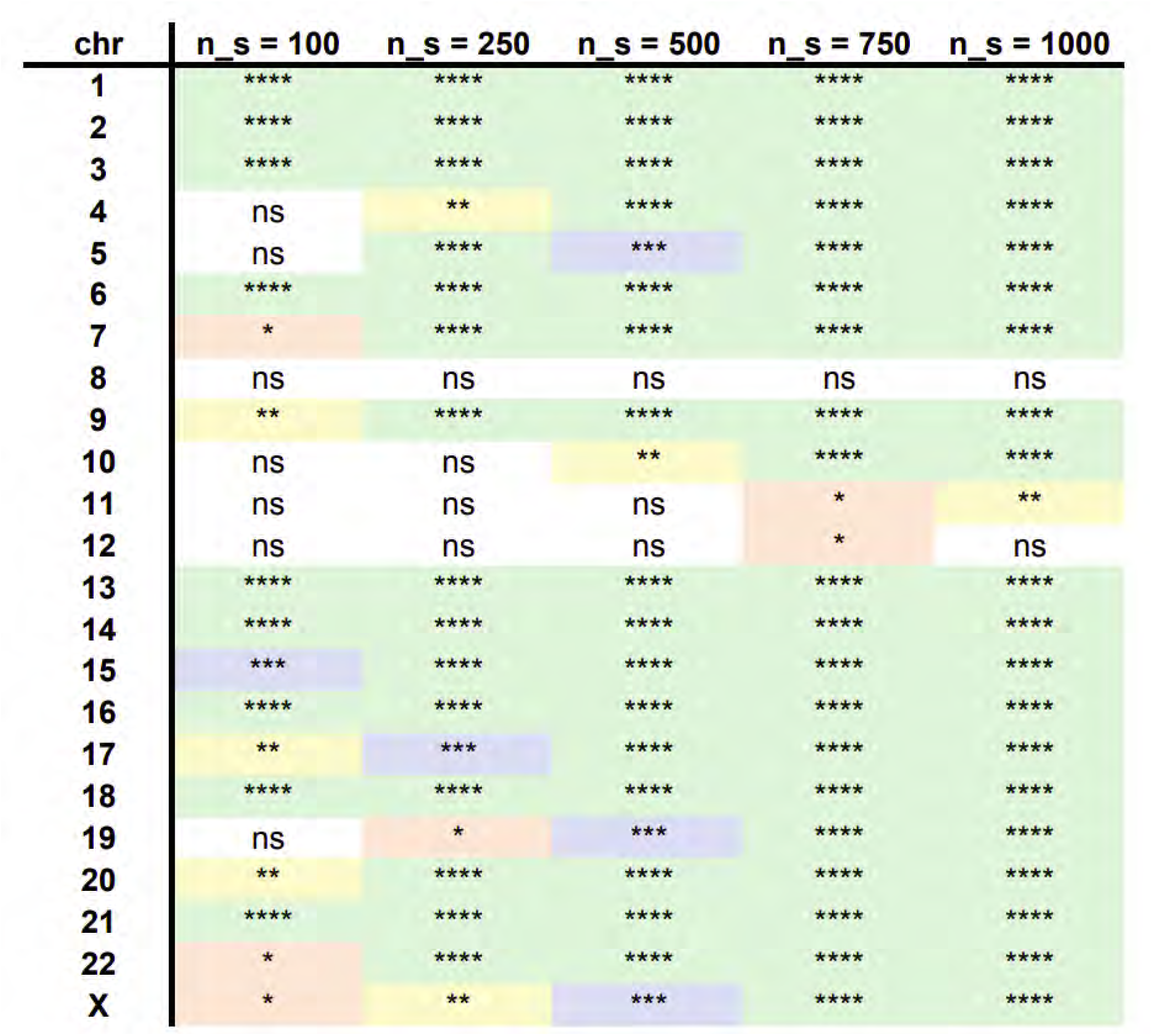
Statistical comparisons of the CT distance profiles obtained from network modeling for GM12878 and BJ1-hTERT for different sample sizes *n*_*s*_ (number of network structures generated from Hi-C data) using the two-sided Mann-Whitney U test. The color values and the asterisk marks represent the level of significance of the BJ1-hTERT vs. GM12878 distribution difference. Green - P-value ≤0.0001 - ****; Blue - P-value ≤ 0.001 - ***; Yellow - P-value ≤0.01 - **; Red - P-value ≤0.05 - *; White - P-value *>* 0.05-non significant (ns)

**Supplementary Figure 10.**
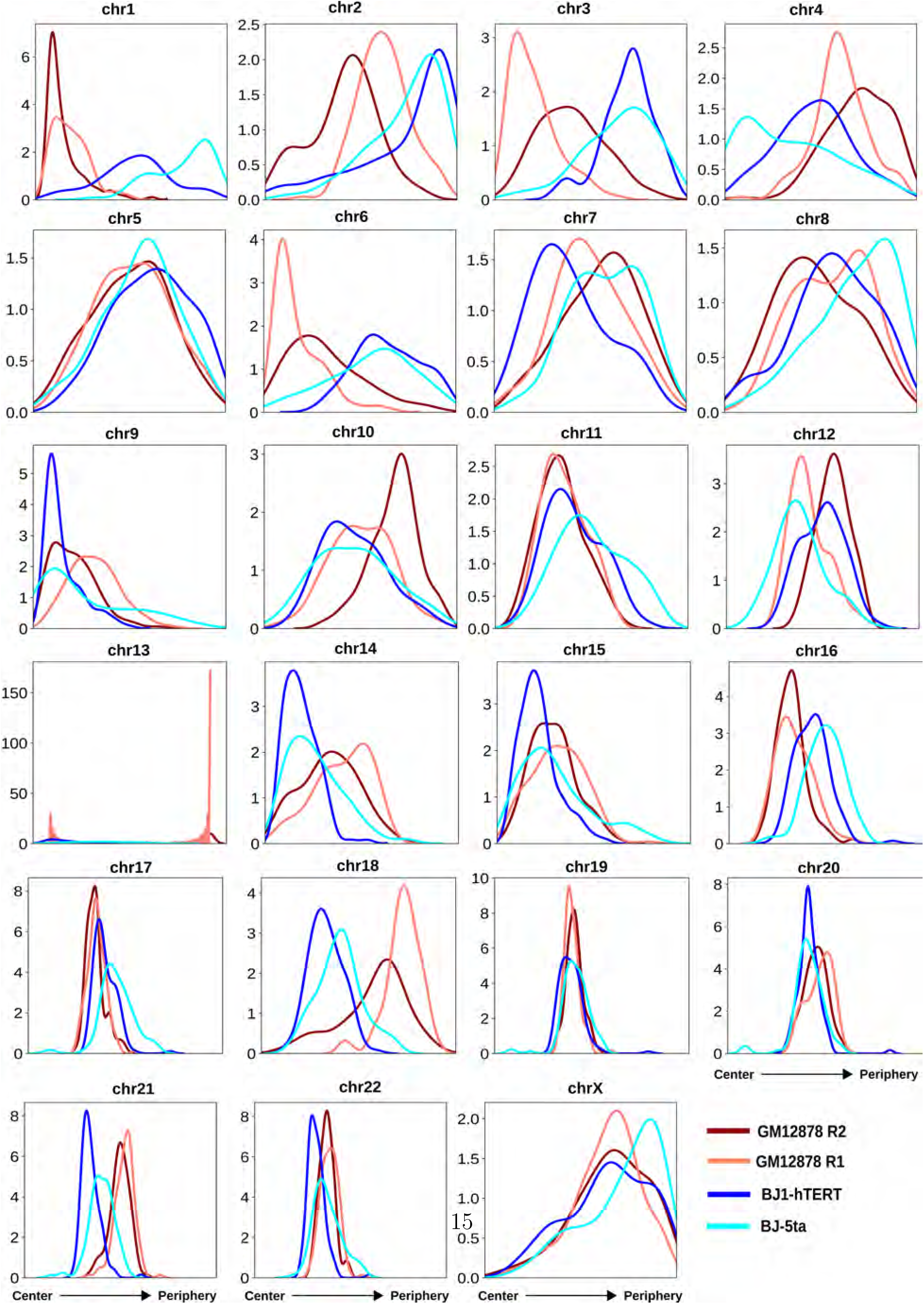
The radial distance profiles of 23 CTs for GM12878 R1, GM12878 R2, BJ1-hTERT and BJ-5ta obtained from the respective selected model clusters based on inferred CT distribution types (# of models in the selected clusters: for GM12878 R2 - 109, GM12878 R1 - 142, BJ1-hTERT - 122, and BJ-5ta - 115)

**Supplementary Figure 11.**
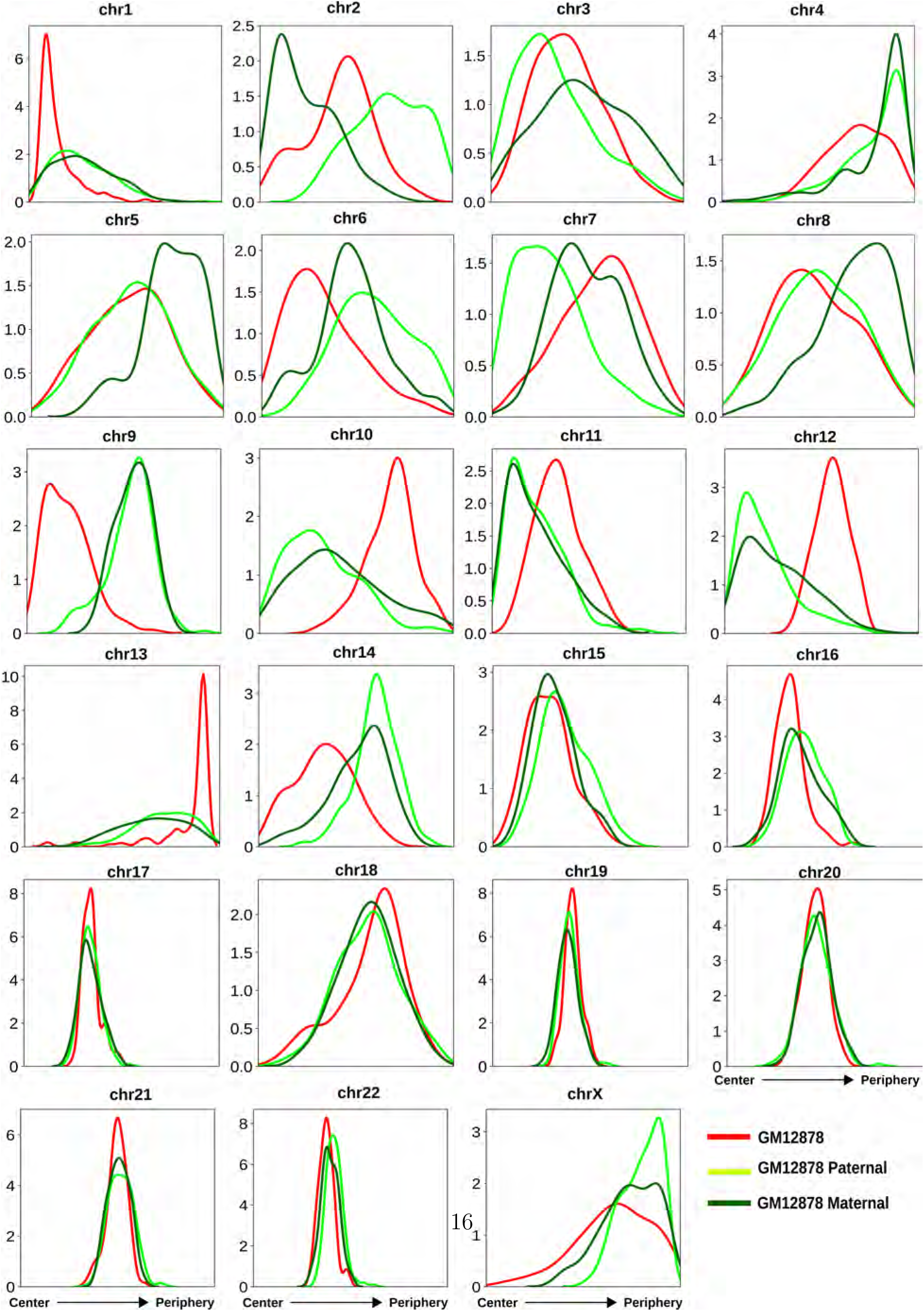
The radial distance profiles of 23 CTs for GM12878 standard, paternal, and maternal copies obtained from the respective selected model clusters based on inferred CT distribution types (# of models in the selected clusters: for GM12878 standard - 109, paternal copy - 194, and maternal copy - 148)

